# Medullary kappa-opioid receptor neurons inhibit pain and itch through a descending circuit

**DOI:** 10.1101/2022.02.08.479623

**Authors:** Eileen Nguyen, Kelly M. Smith, Nathan Cramer, Ruby A. Holland, Isabel H. Bleimeister, Krystal Flores-Felix, Hanna Silberberg, Asaf Keller, Claire E. Le Pichon, Sarah E. Ross

## Abstract

In perilous and stressful situations, the ability to suppress pain can be critical for survival. The rostral ventromedial medulla (RVM) contains neurons that robustly inhibit nociception at the level of the spinal cord through a top-down modulatory pathway. Although much is known about the role of the RVM in the inhibition of pain, the precise ability to directly manipulate pain-inhibitory neurons in the RVM has never been achieved. We now expose a cellular circuit that inhibits nociception and itch in mice. Through a combination of molecular, tracing, and behavioral approaches, we found that RVM neurons containing the kappa-opioid receptor (KOR) inhibit itch and nociception. With chemogenetic inhibition, we uncovered that these neurons are required for stress-induced analgesia. Using intersectional chemogenetic and pharmacological approaches, we determined that RVM^KOR^ neurons inhibit nociception and itch through a descending circuit. Lastly, we identified a dynorphinergic pathway arising from the periaqueductal gray (PAG) that modulates nociception within the RVM. These discoveries highlight a distinct population of RVM neurons capable of broadly and robustly inhibiting itch and nociception.

## Introduction

The brain exerts powerful control over nociceptive processing at the level of the spinal cord. This top-down control occurs through a mechanism known as the descending modulation of pain.^1–3^ Critically, the rostral ventromedial medulla (RVM), represents one final common node within the brain that directly modulates the activity of spinal neurons involved in pain transmission ^4^. Placebo and stress have also been shown to alter the perception of pain through the endogenous opioidergic pain modulatory system, centered upon the RVM.^5–8^ Elucidating how neurons within the RVM can modulate pain presents unique opportunities to harness the brain’s natural pain-inhibitory system to potentially treat chronic pain disorders, thereby avoiding the unwanted and even harmful side effects of existing pharmacological approaches.

Neurons in the RVM have been identified through extensive electrophysiological and pharmacological studies.^4,9^ For example, single unit recordings have provided significant insight into three types of neurons in the RVM: ON, OFF, and NEUTRAL cells based on their firing responses to noxious stimulation and sensitivity to pharmacological agents.^4,10,11^ ON cells are active during noxious stimulation, are inhibited by morphine, and have been proposed to facilitate nociception, whereas OFF cells are inactive or exhibit silent ongoing activity during noxious stimulation, are excited by morphine, and are thought to inhibit nociception.^4,12–14^ Although these studies have established a robust framework by which to classify RVM neurons, many were conducted in anesthetized rodents and the molecular identities of ON and OFF cells remain unclear.

Putative markers that identify subsets of neurons in the RVM have been uncovered, including serotonergic neurons (5HT; marked by *Tph2*), Gad65 (Gad2), neuronal nitric oxide synthase (nNOS; *Nos1*), substance P (*Tac1*) and the kappa-opioid receptor (KOR; *Oprk1*).^15–27^ RVM nNOS neurons are thought to mediate antinociception through local RVM circuits or through projections to cortical and subcortical areas.^20,21^ Serotonergic neurons have classically been presumed to be NEUTRAL cells^28^, although tryptophan hydroxylase (Tph), the enzyme involved in the synthesis of serotonin, has also been identified in cells responsive to mu-agonists, suggesting it may overlap with putative ON cells.^29^ Consistent with this view, optogenetic manipulations of neurons containing Tph have recently revealed their role in facilitating mechanical and thermal hypersensitivity.^15^

The role of inhibitory RVM neurons is also controversial. One study uncovered that RVM^Vgat^ (GABAergic) neurons facilitate mechanical, but not heat, nociception.^19^ However, a different study found that RVM^Gad2^ neurons (which comprise a subpopulation of GABAergic neurons) were involved in the inhibition of heat nociception.^18^ These discordant findings may reflect the fact that GABAergic RVM neurons comprise numerous cell types, and different Cre alleles likely target somewhat different subsets. Indeed, both ON and OFF cells are thought to be GABAergic and GAD+ immunoreactivity has even been identified in some NEUTRAL cells.^30^ In addition to these apparent discrepancies, the role of RVM neurons, including the Gad2/Vgat populations, in the modulation of itch has not been tested.

Previous work has highlighted the existence of OFF cells in the RVM that robustly inhibit nociception^4,12–14^ yet it has been difficult to identify distinct cell types. One candidate population of OFF cells in the RVM includes neurons expressing the kappa-opioid receptor (KOR).^23–25^Microinjections of kappa agonists into the RVM have been shown to elicit mechanical hypersensitivity following injury and attenuate morphine-induced analgesia.^23–25^ These studies, therefore, suggest that RVM^KOR^ neurons participate in the inhibition of nociception. However, this idea remains controversial; microinjections of kappa agonists into the RVM have also resulted in elevations of withdrawal thresholds, which are proposed to occur through the presynaptic inhibition of excitatory inputs onto ON cells^31,32^, thereby resulting in an anti-nociceptive response. One explanation for the observed discrepancies may be due to the site at which kappa agonists exert their action, which could include RVM^KOR^ cell bodies as well as KOR-expressing presynaptic terminals within the RVM. Thus, one advantage of using genetic tools is the ability to manipulate neurons directly, which may shed light on the precise role of RVM^KOR^ neurons.

Here, we screened five putative markers of RVM neurons and discovered the dramatic impact of RVM^KOR^ neurons in the modulation of itch and nociception. In particular, we identified a population of spinally-projecting, GABAergic RVM^KOR^ neurons that inhibit nociceptive thresholds, itch, and pain behaviors following acute and chronic injury. We also determined that RVM^KOR^ neurons tonically inhibit nociception that was unmasked following pharmacological and chemogenetic inhibition. Furthermore, we found that RVM^KOR^ neurons were required for the behavioral expression of stress-induced analgesia. Lastly, we identified that dynorphin signaling from the PAG to the RVM bidirectionally modulates nociception. These findings emphasize the role of the endogenous pain modulatory pathway, centered on RVM^KOR^ neurons, that robustly inhibits nociception and itch.

## Materials and Methods

### Mice

All animals were of the C57Bl/6 background. The studies were performed in both male and female mice 8-10 weeks of age. Even numbers of male and female mice were used for all experiments and no clear sex differences were observed so data were pooled. Mice were given free access to food and water and housed under standard laboratory conditions. The use of animals was approved by the Institutional Animal Care and Use Committee of the University of Pittsburgh.

### Pharmacologic agents

Clozapine-N-oxide (Tocris) was dissolved in PBS and administered intraperitoneally (5 mg/kg), IT (50 μg/kg), or locally via cannula injection (1 mmol in 300 nL). Salvinorin B (SalB; Tocris) was dissolved in DMSO and administered subcutaneously (10 mg/kg). U69,593 (Sigma; 40 ng in 0.25 μL) was dissolved in 45% (2-Hydroxypropyl)-β-cyclodextrin (HBC; Sigma). Morphine sulfate (Henry Schein; 1 μg in 0.25 μL) was diluted in sterile saline. nor-Binaltorphimine dihydrochloride (norBNI; Sigma, 100 ng in 250 nL) was dissolved in sterile saline. U69,593, morphine, and norBNI were microinjected into the RVM. Capsaicin (0.1%) in 10% EtOH in PBS was injected 10 μL into the plantar hindpaw. Chloroquine diphosphate salt (Sigma) was dissolved in physiological saline (100 μg in 10 μL) and administered intradermally. nNOS^CreER^ animals received intraperitoneal (75 mg/kg) tamoxifen (Sigma) dissolved in corn oil over 5 days.

### Intradermal and intrathecal injections

For intradermal injection of chloroquine, hair was removed at least 24 h before the experiment. Chloroquine (100 μg in 10 μL) was administered into the nape of the neck or calf, which could be subsequently visualized by the formation of a small bubble under the skin. For intrathecal injections, hair was clipped from the back of each mouse at least 24 h before the experiment. All intrathecal injections were delivered in a total volume of 10 μL using a 30-gauge needle attached to a luer-tip 25 μL Hamilton syringe. The needle was inserted into the tissue at a 45° angle and through the fifth intervertebral space (L5 – L6). Solution was injected at a rate of 1 μL/s. The needle was held in position for 10 s and removed slowly to avoid any outflow of the solution. Only mice that exhibited a reflexive flick of the tail following puncture of the dura were included in our behavioral analysis. These procedures were performed in awake mice.

### Intraspinal injections

Mice were anesthetized with 100 mg/kg ketamine and 10 mg/kg xylazine. An incision was made at the spinal cord level corresponding to L4-6 dermatome. The intrathecal space was exposed, and two injections of approximately 300 nL of virus was infused 300 mm below the surface of the spinal cord at 5 nL/s via glass pipette through the intrathecal space corresponding to L4-6 of the spinal cord. The same approach was used to target the cervical spinal cord. The glass pipette was left in place for an additional 5 min before withdrawal. The incision was closed with 6-0 vicryl suture. Ketofen was delivered (IP 10 mg/kg) and mice were allowed to recover over a heat pad.

### Stereotaxic injections and cannula implantation

Animals were anesthetized with 2% isoflurane and placed in a stereotaxic head frame. A drill bit (MA Ford, #87) was used to create a burr hole and a custom-made metal needle (33 gauge) loaded with virus was subsequently inserted through the hole to the injection site. Virus was infused at a rate of 100 nL/min using a Hamilton syringe with a microsyringe pump (World Precision Instruments). Mice received 250-500 nL of virus. The injection needle was left in place for an additional 15 min and then slowly withdrawn over another 15 min. Injections and cannula implantations were performed at the following coordinates for each brain region: RVM: AP -5.80 mm, ML 0.00 mm, DV -6.00 and PAG: AP -4.70 mm, ML ± 0.74 mm, DV: -2.75. For implantation of guide cannulas (P1 Technologies), implants were slowly lowered 0.300-0.500 mm above the site of injection and secured to the skull with a thin layer of Vetbond (3M) and dental cement. The incision was closed using Vetbond and animals were given ketofen (IP 10 mg/kg) and allowed to recover over a heat pad. Mice were given 4 weeks to recover prior to experimentation.

### Spared nerve injury

For spared nerve injury surgery (SNI), the sural and superficial peroneal branches of the sciatic nerve were ligated (size 6.0 suture thread) and transected, leaving the tibial nerve intact. The muscle tissue overlying the injury was gently placed back together and the skin sutured. Animals were tested 3 weeks post-injury.

### Capsaicin-induced injury

Animals received 10 μL of intraplantar capsaicin (0.1% w/v in 10% ethanol diluted in saline) and the total licks and time spent licking the injured paw was quantified over 20 minutes.

### Behavior

All assays were performed in the Pittsburgh Phenotyping Core and scored by an experimenter blind to treatment.

### Observation of scratching behavior

Scratching behavior was observed using a previously reported method.^33^ On the testing day, the mice were individually placed in the observation cage (12×9×14 cm) to permit acclimation for 30 min. Scratching behavior was videotaped for 30 min after administration of chloroquine. The temporal and total numbers of scratch bouts by the hind paws at various body sites during this period were counted. For the experiments targeting the lumbar spinal cord, the amount of time spent biting following intradermal chloroquine was summated.

### Hargreaves testing

Animals were acclimated on a glass plate held at 30°C (Model 390 Series 8, IITC Life Science Inc.). A radiant heat source was applied to the hindpaw and latency to paw withdrawal was recorded.^34^ Two trials were conducted on each paw, with at least 5 min between testing the opposite paw and at least 10 min between testing the same paw. To avoid tissue damage, a cut off latency of 20 sec was set. Values from both paws were averaged to determine withdrawal latency.

### Stress-induced analgesia (forced swim test)

Mice were placed in a water bath maintained at 30 C and forced to swim for 3 min. After the swim, mice were returned to their home cages and tested 30 min later for stress-induced analgesia to hotplate and capsaicin (described further in the methods section).

### Hotplate

Twenty minutes after the forced swim test, mice were placed on a 55 C hotplate. The latency to first nocifensive response (lick or jump) and total number of licks over a 60 s period were measured. Values were averaged across two trials for each mouse.

### von Frey testing

Mechanical sensitivity was measured using the Chaplan up-down method of the von Frey test.^35^ Calibrated von Frey filaments (North Coast Medical Inc.) were applied to the plantar surface of the hindpaw. Lifting, shaking, and licking were scored as positive responses to von Frey stimulation. Average responses were obtained from each hindpaw, with 3 min between trials on opposite paws, and 5 min between trials on the same paw.

### Cold plantar assay

Paw withdrawal to cold sensitivity was measured as previously described.^36,37^ Briefly, animals were acclimated to a 1/4” glass plate and crushed dry ice was packed into a modified 3 mL syringe and used as a probe. The dry ice probe was applied to the glass beneath the plantar hindpaw and the latency to withdrawal was recorded. Two trials were conducted on each hindpaw, with 3 min between trials on opposite paws, and 5 min between trials on the same paw. A cut off latency of 20 s was used to prevent tissue damage. Withdrawal latencies for each paw were determined by averaging values across 2 trials per paw.

### Tail-flick assay

Tails were immersed 3 cm into a water bath at 48°C, and the latency to tail-flick was measured three times with a 1 min interval between trials.

### Rotarod

Coordination and balance were determined using an accelerating rotarod. A session involving 120 s on a non-accelerating rotarod was prior to experimentation. Five consecutive trials were performed while the rotarod accelerated from 4 to 40 rpm in 30 s increments. The latency to fall off (s) was recorded for each trial.

### Open field activity

Spontaneous activity in the open field was conducted over 30 min in an automated Versamax Legacy open field apparatus for mice (Omnitech Electronics Incorporated, Columbus, OH). Distance traveled was measured by infrared photobeams located around the perimeter of the arenas interfaced to a computer running Fusion v.6 for Versamax software (Omnitech Electronics Incorporated) which monitored the location and activity of the mouse during testing. Activity plots were generated using the Fusion Locomotor Activity Plotter analyses module (Omnitech Electronics Incorporated). Mice were placed into the open field 30 min post CNO injection.

### Acoustic startle

Prepulse inhibition of acoustic startle was measured by placing mice into a soundproof chamber (Kinder Scientific Startle Monitor). After 5 min of white noise acclimation, mice were exposed to randomized trials of 500 ms exposures to 65 or 115 decibel sound pressure level (dB SPL) with a 500 ms intertrial interval. Trials were repeated between seven and eight times for each mouse in a randomized order. Startle was measured as the maximum force in Newtons (N) and the average response across the trial repetitions for each mouse was used for data analysis.

### RNAscope in situ hybridization

Multiplex fluorescent in situ hybridization (FISH) was performed according to the manufacturer’s instructions (Advanced Cell Diagnostics #320850). Briefly, 14 um-thick fresh-frozen sections containing the RVM were fixed in 4% paraformaldehyde, dehydrated, treated with protease for 15 min, and hybridized with gene-specific probes to mouse. Probes were used to detect tdTomato-C2 (#317041-C2), mCherry-C2 (#431201), Mm-Oprm1-C1 (#315841), Mm-Oprk1-C1 (#316111), Mm-Nos1-C2 (#437651-C2), Mm-Tac1-C1 (#517971), Mm-Tph2-C2 (#318691-C2), Mm-Fos-C3 (#498401-C3), Mm-Pdyn-C2 (#31877), Mm-Slc32a1-C3 (#319191), and Mm-Slc17a6-C3 (#319171). DAPI (#320858) was used to visualize nuclei. 3-plex positive (#320881) and negative (#320871) control probes were tested. Three to four z-stacked sections were quantified for a given mouse, and 3-4 mice were used per experiment.

### Immunohistochemistry

Mice were anesthetized with an intraperitoneal injection of urethane, transcardially perfused, and post-fixed at least 4 h in 4% paraformaldehyde. 40 μm and 25 μm thick RVM and spinal cord sections were collected on a cryostat and slide-mounted for immunohistochemistry. Sections were blocked at room temperature for 2 h in a 5% donkey serum, 0.2% triton, in phosphate buffered saline. Primary antisera was incubated for 14 h overnight at 4° C: rabbit anti-RFP (1:1K), chicken anti-GFP (1:1K), and mouse anti-NeuN (1:500). Sections were subsequently washed three times for 20 min in wash buffer (0.2% triton, in PBS) and incubated in secondary antibodies (Life Technologies, 1:500) at room temperature for 2 h. Sections were then incubated in Hoechst (ThermoFisher, 1:10K) for 1 min and washed 3 times for 15 min in wash buffer, mounted and cover slipped.

### Fos experiments

Morphine sulfate (1 μg in 0.25 μL) was microinjected into the RVM. Twenty minutes thereafter, brain samples were collected and processed for FISH. Spinal cords were harvested 20 min and 90 min following CNO administration for FISH and immunohistochemistry, respectively.

### Image acquisition and quantification

Full-tissue thickness sections were imaged using either an Olympus BX53 fluorescent microscope with UPlanSApo 4x, 10x, or 20x objectives or a Nikon A1R confocal microscope with 20X or 60X objectives. All images were quantified and analyzed using ImageJ. To quantify images in RNAscope in situ hybridization experiments, confocal images of tissue samples (3-4 sections per mouse over 3-4 mice) were imaged and only cells whose nuclei were clearly visible by DAPI staining and exhibited fluorescent signal were counted. To quantify spinal fluorescence, we measured the mean intensity of fluorescence within the spinal cord (both dorsal and ventral horns) using ImageJ.

### Tissue clearing by iDisco and light sheet imaging

The iDisco+ protocol was followed as previously described.^38^ Whole brains and brainstem-spinal cord samples were immunolabeled with rabbit anti-RFP (Rockland #600-401-379) at 1:200 during a 7-day incubation at 37 °C. Secondary antibody AlexaFluor 647-highly cross-adsorbed donkey anti-rabbit (Thermo Fisher, Cat #A-31573) was used at 1:400. iDisco-processed samples were imaged using light sheet microscopy (Miltenyi Biotec Ultramicroscope II, with the 1x and 4x objectives). All images were acquired with illumination from a single side (three light sheets) with dynamic focusing (10 positions) enabled. The ImSpector software ‘Blend’ algorithm was used for dynamic focus processing. Images were stitched using Arivis Vision4D software.

### Spinal cord slice preparation

Four weeks after surgery, spinal cord slices were prepared using methods described previously (Graham et al 2003 and Graham et al 2007). In brief, mice were deeply anaesthetized with a ketamine/xylazine cocktail (90 mg/kg, 10 mg/kg respectively) and decapitated. The spinal cord was dissected in ice cold NMDG aCSF containing (in mM): 93 NMDG, 2.5 KCl, 1.2 NaH_2_PO_4_, 30 NaHCO_3_, 20 HEPES, 25 Glucose, 5 sodium ascorbate, 2 thiourea, 3 sodium pyruvate, 10 MgSO_4_.7H_2_O, 0.5 CaCl_2_.2H_2_O, pH 7.3-7.4 with HCl. Parasagittal slices (200µM) from lumbar spinal cord were made using a vibrating microtome (Leica VT1200s). Slices were transferred to oxygenated, room temperature aCSF containing (in mM): 124 NaCl, 2.5 KCl, 1.2 NaH_2_PO_4_, 24 NaHCO_3_, 5 HEPES, 12.5 Glucose, 2 MgSO_4_.7H_2_O, 2 CaCl_2_.2H_2_O, pH 7.3-7.4, and stored in an immersion incubation chamber for at least 1 h before recording.

### RVM slice preparation and electrophysiology

Acute brain slices were generated from KOR^Cre^ mice in which spinally projecting RVM^KOR^ neurons had been labeled by injection of AAVr-FLEX-tdTomato into the lumbar spinal. Under ketamine/xylazine anesthesia, mice were perfused transcardially with ice-cold NMDG aCSF and horizontal sections of the brainstem, 300 µm thick, were cut on a Leica VT1200s vibratome. Slices were allowed to recover for 7 minutes in NMDG aCSF at 33°C and then transferred to normal aCSF to recover for 1 hour at room temperature before recording. Whole cell recordings were obtained from fluorescent cells with borosilicate glass pipettes containing (in mM): 70 K-Gluconate, 60 KCl, 10 HEPES, 1 MgCl_2_, 0.5 EGTA, 2.5 Mg-ATP, 0.2 GTP-Tris, with pH of 7.3 and 285 mOsm. Miniature inhibitory postsynaptic currents (mIPSCs) were isolated with bath application of (in µM): 0.5 TTX, 20 CNQX and 50 APV. The µ-opioid receptor agonist DAMGO (abcam) was bath applied at 10 nM. Data were acquired using MultiClamp 700B amplifier and Axon Digidata 1550B and stored on a computer running Clampex software (Molecular Devices, San Jose CA). Spontaneous inhibitory postsynaptic currents (sIPSC) were isolated with MiniAnalysis (Synaptosoft) and analyzed in Prism (GraphPad, San Diego CA).

### Patch clamp electrophysiology

Slices were transferred to the recording chamber perfused with oxygenated, room temperature aCSF. Neurons were visualized using a 40x water immersion objective and IR-DIC optics. Recordings were then made from randomly selected neurons within or dorsal to the substantia gelatinosa, easily visualized under brightfield illumination by its translucent appearance. Patch pipettes (6-9 MΩ) were filled with a CsCl-based internal solution containing (in mM): 130 CsCl, 1 MgCl_2_.6H_2_O, 10 HEPES, 2 Mg_2_ATP, 0.3 Na_3_GTP, 10 EGTA, pH 7.3 with CsOH. Data was captured using a Multiclamp 200B amplifier (Molecular Devices), digitized with an ITC-18 (Instrutech) and stored using Axograph X software (Molecular Devices).

Photostimulation of KOR^Cre^ fibers originating from the RVM was achieved using a high intensity LED light source evoked by the data acquisition program (Axograph) delivered through the microscopes optical path. Photostimulation (488 nm) of increasing duration was delivered at a rate of 0.5 Hz (1ms, 5 ms, 50 ms, 500 ms and 1 s). Responses of dorsal horn neurons to photostimulation was recorded in voltage clamp with a holding potential of -70 mV. CNQX (10 µM, Sigma-Aldrich) was included in all recordings to ensure capture of inhibitory events only. Each light stimulation protocol was repeated 10 times per cell with a 7 s gap between each sweep. In a subset of recordings, bicuculline (10 µM, Sigma-Aldrich) was added to the bath solution.

### Electrophysiology data analysis

All data was analyzed offline using Axograph software. Connections between KOR cre fibers originating from the RVM and spinal cord neurons were counted if optically evoked currents were present in at least 50% of trials. Optically evoked inhibitory post synaptic currents (oIPSCs) for each stimulation duration were averaged and maximum amplitude and latency from stimulus was measured. oIPSC latency was also measured from individual trials to determine jitter (defined as 2 x standard deviation) for each stimulus duration.

### Statistical analysis

For behavioral, FISH, and neurochemical experiments, statistical analyses were performed using GraphPad Prism 8. Values are presented as mean ± SEM. Statistical significance was assessed using tests indicated in applicable figure legends. Significance was indicated by p < 0.05. Sample sizes were based on pilot data and are similar to those typically used in the field.

## Data availability

Further information and requests for resources and reagents should be directed to and will be fulfilled by the corresponding authors and Lead Contact, Dr. Sarah E. Ross.

## Results

### Evaluation of RVM cell types in the descending modulation of nociception

Neurons in the RVM have been classified functionally as ON, OFF, or NEUTRAL cells based on their response properties. They have also been classified neurochemically based on their expression of neurotransmitters and other markers, such as GABA, serotonin, nNOS, Tac1, and KOR. However, we do not yet have a clear understanding of which neurochemical classes correspond to which functional classes. To address this gap, we began by characterizing the neuronal diversity of the RVM through multiplex fluorescent in-situ hybridization (FISH) (Fig. 1A, Supplementary Fig. 1A). We found that 37.3 ± 17.1% of serotonergic/*Tph2* and 74.8 ± 3.0% of KOR/*Oprk1* were GABAergic (based on their expression of *Slc32a1*). *Tac1* neurons were glutamatergic (based on co-expression of *Slc17a6*; 41.3 ± 14.2%), and Nos1 neurons were both GABAergic and glutamatergic (16.6 ± 7.9% and 49.7 ± 14.8%, respectively). Thus, the markers that we analyzed exhibited mixed neurotransmitter phenotypes, underscoring the complexity of neurons within the RVM (Fig. 1A, Supplementary Fig. 1A).

**Figure 1.**
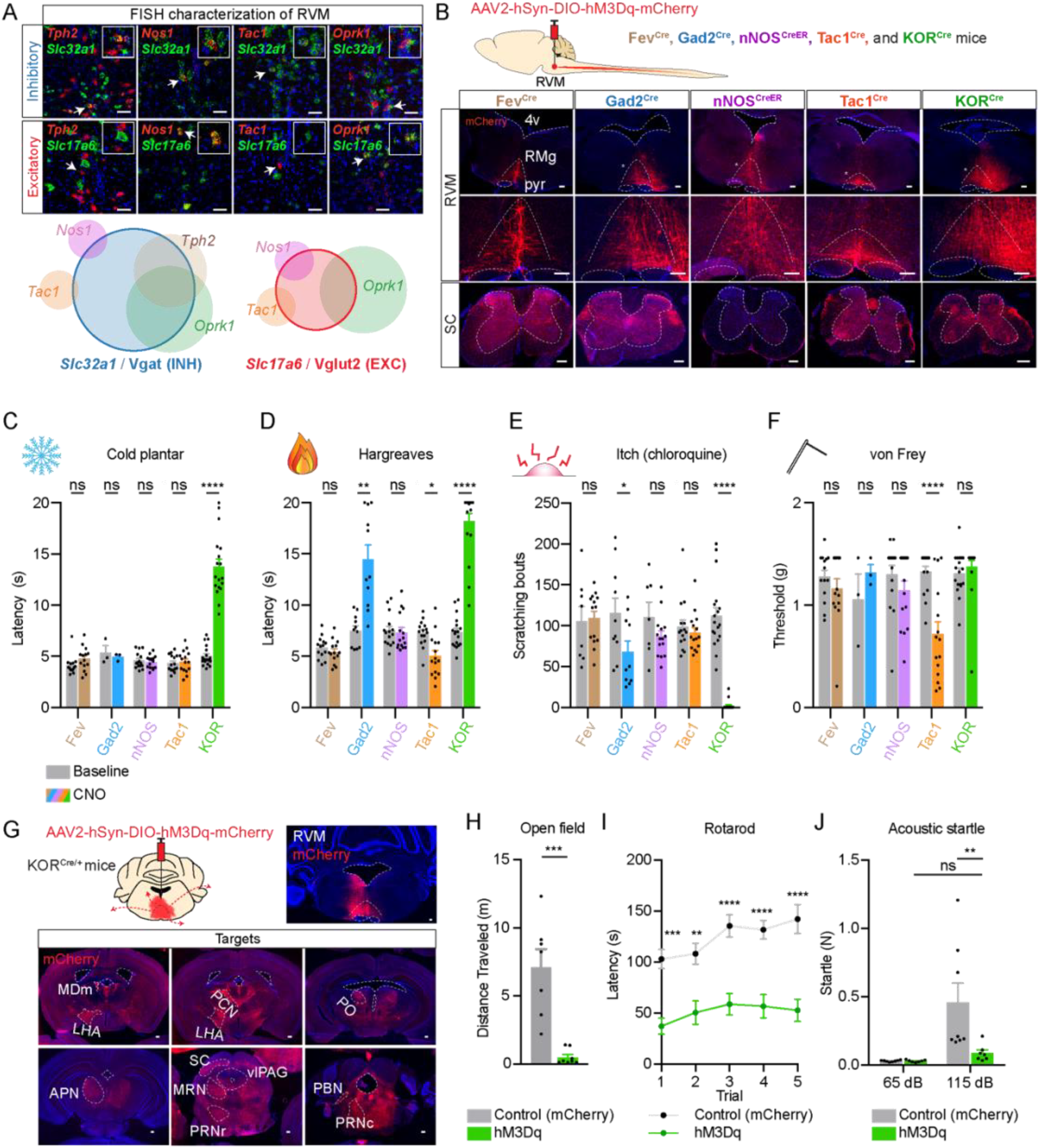
Identification and manipulation of diverse RVM neuronal cell types. **(A)** Characterization of neuronal populations in the rostral ventromedial medulla (RVM) using fluorescent in-situ hybridization (FISH). Intersecting Venn diagrams indicate the relative abundance of neurons expressing particular markers and their overlap. Quantifications are provided in Supplementary Figure 1B. **(B)** Experimental approach to deliver AAV2-hSyn-DIO-hM3Dq-mCherry to visualize RVM neurons and their spinal projections across Fev^Cre^ (serotonin), Gad2^Cre^, nNos^CreER^, Tac1^Cre^, and KOR^Cre^ alleles. SC: spinal cord, 4v: fourth ventricle, RMg: raphe magnus, pyr: pyramidal tract. Scale bar = 100 μm. **(C-F)** Effect of chemogenetic activation of Fev^Cre^, Gad2^Cre^, nNos^CreER^, Tac1^Cre^, and KOR^Cre^ neurons on sensory behaviors. Asterisks indicate the results of two-way repeated measures ANOVA, with Bonferroni’s multiple comparisons test, between baseline and following intraperitoneal (IP) CNO (5 mg/kg). Data are mean + SEM with dots representing individual mice. (ns p>0.05, * p<0.05, ** p<0.01, **** p<0.0001). **(C)** Cold plantar assay. Number of mice for each allele: Fev^Cre^ (n = 14), Gad2^Cre^ (n = 3), nNos^CreER^ (n = 15), Tac1^Cre^ (n = 15), KOR^Cre^ (n = 18). **(D)** Hargreaves assay. Number of mice for each allele: Fev^Cre^ (n = 14), Gad2^Cre^ (n = 11), nNos^CreER^ (n = 15), Tac1^Cre^ (n = 15), KOR^Cre^ (n = 18). **(E)** Itch assay (chloroquine-evoked itch). Number of mice for each allele: Fev^Cre^ (n = 8), Gad2^Cre^ (n = 10), nNos^CreER^ (n = 7), Tac1^Cre^ (n = 15), KOR^Cre^ (n = 18). **(F)** von Frey assay. Number of mice for each allele: Fev^Cre^ (n = 14), Gad2^Cre^ (n = 3), nNos^CreER^ (n= 15), Tac1^Cre^ (n = 15), KOR^Cre^ (n = 18). **(G)** Delivery of AAV2-hSyn-DIO-hM3Dq-mCherry to KOR^Cre^ mice results in infection of neurons within the RVM as well as beyond the RVM in the brainstem. Representative images of projections to supraspinal areas are shown. MDm: mediodorsal nucleus of the thalamus, medial part, LHA: lateral hypothalamic area, PCN: paracentral nucleus, PO: posterior complex of the thalamus, APN: anterior pretectal nucleus, SC: superior colliculus, vlPAG: ventrolateral periaqueductal gray, MRN: midbrain reticular nucleus, PRNr: pontine reticular nucleus, PBN: parabrachial nucleus, PPNc: pontine reticular nucleus, caudal part. Scale bar = 50 μm. **(H-J)** Effect of activation of RVM^KOR^ neurons using AAV2-hSyn-DIO-hM3Dq-mCherry in KOR^Cre^ mice on locomotor and acoustic startle assays. Control animals received AAV2-hSyn-DIO-mCherry. Data are mean + SEM with dots representing individual mice (ns p>0.05, ** p<0.01, *** p<0.001, **** p<0.0001). **(H)** Open field assay (n = 7 controls, n = 8 hM3Dq mice). Asterisks indicate the results of unpaired t-test (*** p<0.001). **(I)** Rotarod assay. Data are mean ± SEM (n = 8 controls, n = 14 hM3Dq mice). Asterisks indicate the results of two-way repeated measures ANOVA with Bonferroni’s correction. **(J)** Acoustic startle assay (n = 8 controls, n = 7 hM3Dq mice). Asterisks indicate the results of two-way repeated measures ANOVA with Bonferroni’s correction.

To visualize and manipulate these cell groups, we acquired the corresponding Cre strains (serotonin/Fev^Cre^, Gad2^Cre^, nNOS^CreER^, Tac1^Cre^, and KOR^Cre^) and targeted the RVM in these mice through stereotaxic injections of Cre-dependent AAVs encoding hM3Dq-mCherry for activation, or mCherry alone as a control (Fig. 1B, Supplementary Fig. 1B). As expected, based on FISH, numerous neurons in the RVM were targeted with each of the Cre alleles (Fig. 1B). Most of the genetically defined populations exhibited prominent RVM projections to the spinal cord, consistent with the possibility that these cell types play a role in descending modulation (Fig. 1B). Although the nNOS population did not display significant spinal innervation (Fig. 1B), we also tested its possible indirect role on nociception.

Next, we performed a screen to examine whether the activation of RVM neurons targeted by any of these Cre alleles affected somatosensory responses across four major distinct modalities: noxious cold (cold plantar test), noxious heat (Hargreaves test), itch (chloroquine test), and mechanical force (von Frey test) (Fig. 1C-F). Neither activation of Fev^Cre^ nor nNOS^CreER^ neurons had any significant effect in the somatosensory assays tested. In contrast, activation of Gad2^Cre^ neurons significantly increased thermal thresholds, consistent with the previous report that this population is anti-nociceptive.^18^ Activation of Tac1^Cre^ neurons had the opposite effect: it decreased response latencies in the Hargreaves test and thresholds in the von Frey test. Of all the alleles tested, the one with the most prominent phenotypes was the KOR^Cre^ allele. Chemogenetic activation of the KOR^Cre^ population increased response latencies to cold and heat while completely blocking scratching in response to chloroquine (Fig. 1C-E).

Given the dramatic findings with the KOR^Cre^ allele observed in our initial screen, we began analyzing the targeted cells in more detail. Although the viral injections into the KOR^Cre^ mice were generally confined to the RVM and cells just around this region, we observed many projections from mCherry-labeled KOR^Cre^ neurons to supraspinal areas including the lateral hypothalamus, parabrachial nucleus, and superior colliculus (Fig. 1G), suggesting that numerous regions may have been affected by chemogenetic manipulation. The labeling observed throughout the brain suggested that we may have also targeted KOR-expressing neurons in surrounding nuclei.

Moreover, when we performed other behavioral tests using these KOR^Cre^ mice, we found that they showed a generalized lack of responsiveness in multiple assays (Fig. 1H-J). Compared to their control counterparts, the mice in which KOR^Cre^ neurons in the RVM area were chemogenetically activated moved less (open field test), displayed a lack of coordination (decreased rotarod latency) and showed a reduced startle response to loud noises (acoustic startle). These findings, together with the diminished somatosensory responses, raised the possibility that KOR neurons in the RVM as well as neurons in surrounding regions in the brainstem have a profound effect on arousal.

To disambiguate these phenotypes, we decreased the viral titer used in our stereotaxic injections through a serial dilution study (from 8.6×10^10^ to 2.7×10^9^, i.e. 32-fold) (Supplementary Fig. 2B-D). With very dilute titers that were restricted to the RVM, the only projection out of the RVM that was observed in RVM^KOR^ neurons was that directed to the spinal cord (Supplementary Fig. 2D, E). To further visualize that our diluted titer indeed captured only local and spinally-projecting neurons, we used an iDisco clearing approach. We compared labeling of RVM^KOR^ projections throughout the brain and spinal cords of animals receiving viral titers with and without dilution (undiluted titer: 8.6×10^10^; diluted viral titer: 4.3×10^9^, i.e. 20-fold diluted) (Supplementary Fig. 2E, Supplementary Video 1 and 2). Consistent with our histology of coronal sections (Supplementary Fig. 2D, F), our intact iDisco-cleared brainstem-spinal samples revealed that the use of the diluted viral titer resulted in labeling of descending RVM^KOR^ neurons, with limited labeling of supraspinal projections (Supplementary Fig. 2F, Supplementary Video 1 and 2). Importantly, using the lowest viral titers, we were able to see a clear correlation between correct targeting of KOR neurons in the RVM (as evidenced by spinal projections) and the presence of a sensory phenotype upon chemogenetic activation of the infected cells (Supplementary Fig. 2C, D).

Having fine-tuned our viral strategy for more selective targeting of RVM^KOR^ neurons (Fig. 2A), we next characterized these cells in more detail. First, our FISH experiments revealed that mCherry-expressing RVM^KOR^ neurons comprised 88.6 ± 3.6% of *Oprk1* neurons in the RVM and that these neurons were 100% specific to the *Oprk1* population (Fig. 2A). Of note, the *Oprk1* neurons in the RVM were diverse with respect to neurotransmitter, with 75-80% of KOR *(Oprk1)* neurons co-expressing *Slc32a1* (Vgat) and *Gad2*, 25% co-expressing *Slc17a6* (Vglut2), 46% co-expressing *Tph2* (serotonin), and 30% overlapping with *Penk* (Fig. 2B). Moreover, approximately 30% of *Oprk1* neurons co-expressed *Oprm1* (MOR), consistent with prior reports.^30,39^ In contrast, there was no overlap of *Oprk1* with either *Nos1* (nNOS) or *Tac1* (Fig. 2B, Supplementary Fig. 2A).

**Figure 2.**
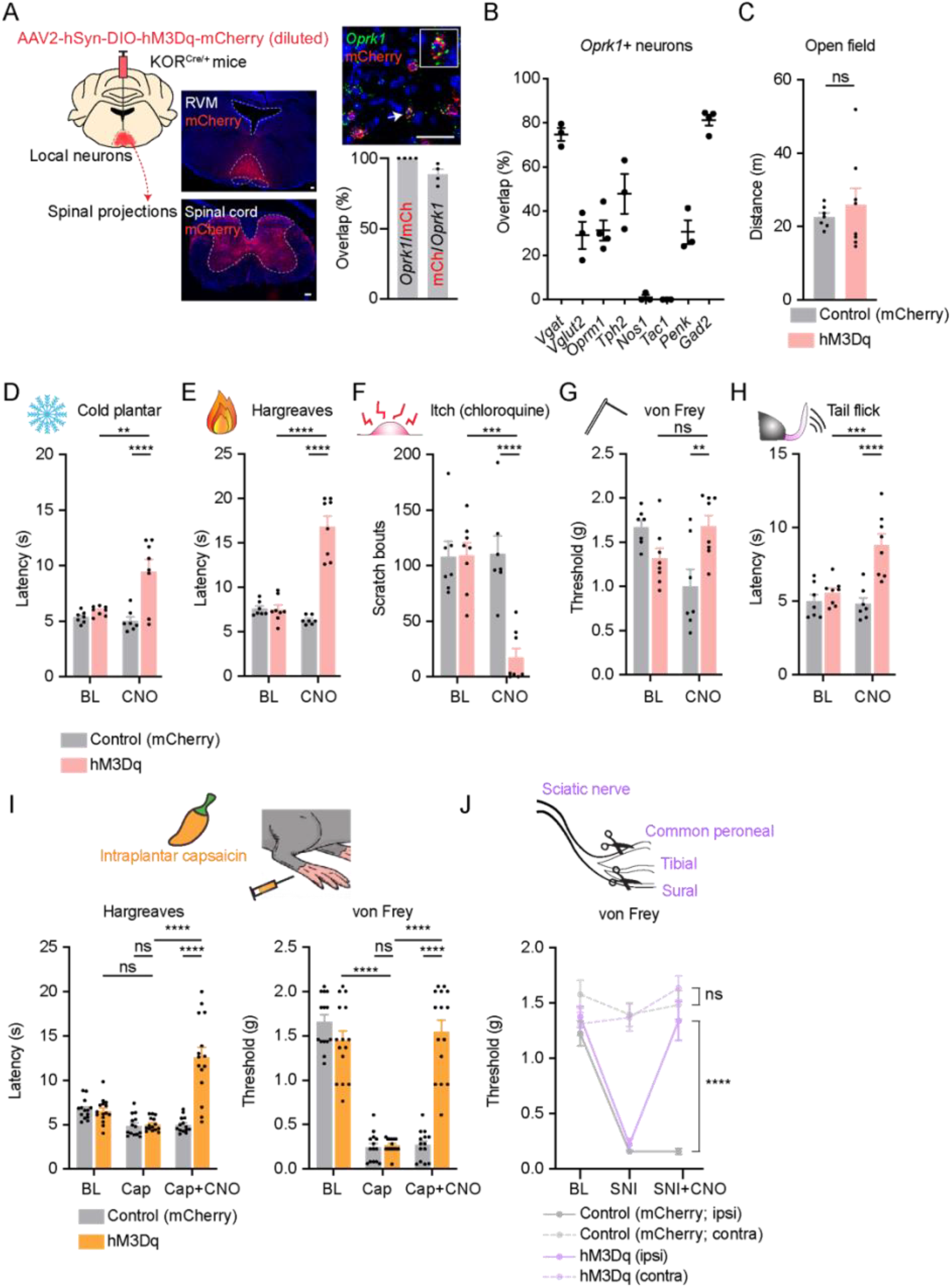
Activation of RVM^KOR^ neurons is analgesic. **(A)** Validation of the diluted viral approach to target RVM^KOR^ neurons using FISH. Data are mean + SEM with dots representing individual mice (n = 4). Scale bar = 50 μm. **(B)** FISH characterization of *Oprk1*-expressing neurons in the RVM relative to other markers of interest. Data are mean ± SEM with dots representing individual mice (n = 3-4 mice per group). **(C)** Effect of chemogenetic activation on locomotor activity in an open field test. Data are mean + SEM with dots representing individual mice (n = 7 controls, n = 8 hM3Dq mice). NS indicates the results of unpaired t-test (ns p>0.05). **(D to H)** Effect of chemogenetic activation of RVM^KOR^ neurons with IP CNO (5 mg/kg) on naive responses to **(D)** cold plantar assay, **(E)** Hargreaves assay, **(F)** chloroquine-evoked itch, **(G)** von Frey assay, and **(H)** tail-flick assay. Data are mean + SEM with dots representing individual mice (n = 7 controls, n = 8 hM3Dq). Asterisks indicate the results of two-way repeated measures ANOVA with Bonferroni’s correction (** p<0.01, *** p<0.001, **** p<0.0001). **(I)** Effect of activation of RVM^KOR^ cells on thermal and mechanical responses with intraplantar capsaicin-induced injury. Data are mean + SEM with dots representing individual mice (n = 14 controls, n = 15 hM3Dq). Asterisks indicate the results of two-way repeated measures ANOVA with Bonferroni’s correction (ns p>0.05, **** p<0.0001). **(J)** Effect of chemogenetic activation of RVM^KOR^ on mechanical hypersensitivity with spared nerve injury. Data are mean ± SEM (n = 14 controls, n = 15 hM3Dq). Asterisks indicate the results of two-way repeated measures ANOVA with Bonferroni’s correction (ns p>0.05, **** p<0.0001).

### RVM^KOR^ neurons inhibit nociceptive thresholds, itch, and acute and chronic pain

Next, we tested the effect of chemogenetically activating RVM^KOR^ neurons. Importantly, with selective targeting of the RVM, mice no longer displayed a generalized defect in arousal as evidenced by normal behavior in the open field test (Fig. 2C). However, in somatosensory assays, chemogenetic activation of RVM^KOR^ neurons produced elevated thresholds to cold, heat and mechanical testing as well as decreased itch behavior (Fig. 2D-G). Since RVM neurons have been traditionally classified based on their firing patterns in relation to the tail-flick response^4,10,11^, we performed this assay as well (Fig. 2H). Activation of RVM^KOR^ neurons likewise caused elevated tail-flick latencies, consistent with the notion that they could be OFF cells. To test the role of RVM^KOR^ neurons in ongoing pain, we analyzed hypersensitivity following intraplantar capsaicin (acute pain) and following nerve injury (chronic pain). Following intraplantar capsaicin, mice developed thermal and mechanical hypersensitivity, and this effect was reversed with activation of RVM^KOR^ neurons (Fig. 2I, Supplementary Fig. 3A-B). Similarly, following spared nerve injury (SNI), both control and hM3Dq groups exhibited pronounced mechanical allodynia, but this hypersensitivity was abrogated following chemogenetic activation of RVM^KOR^ neurons (Fig. 2J). Together, these data suggest that RVM^KOR^ neurons are sufficient for the inhibition of nociception, itch, and the reversal of pain induced by acute (capsaicin) and chronic (SNI) injury.

Microinjection of a kappa agonist into the RVM has previously been shown to block the analgesia driven by mu agonists within other sites of the CNS^23,32^ and can precipitate mechanical allodynia following spinal nerve ligation.^40^ Thus, we sought to determine whether microinjection of a kappa agonist, U69,593, could affect naive responses to nociceptive stimuli (Fig. 3A). When microinjected into the RVM, U69,593 had a pro-nociceptive effect and reduced the latencies to respond to cold and thermal stimuli (Fig. 3B, C) as well as mechanical thresholds (Fig. 3D). Consistent with previous studies^23^, microinjection of U69,593 did not affect latencies in response to tail-flick (Fig. 3E). These data support that, at least for some stimuli, kappa-sensitive neurons in the RVM tonically suppress responses to nociception that may be unmasked with their temporary pharmacological inhibition.

**Figure 3.**
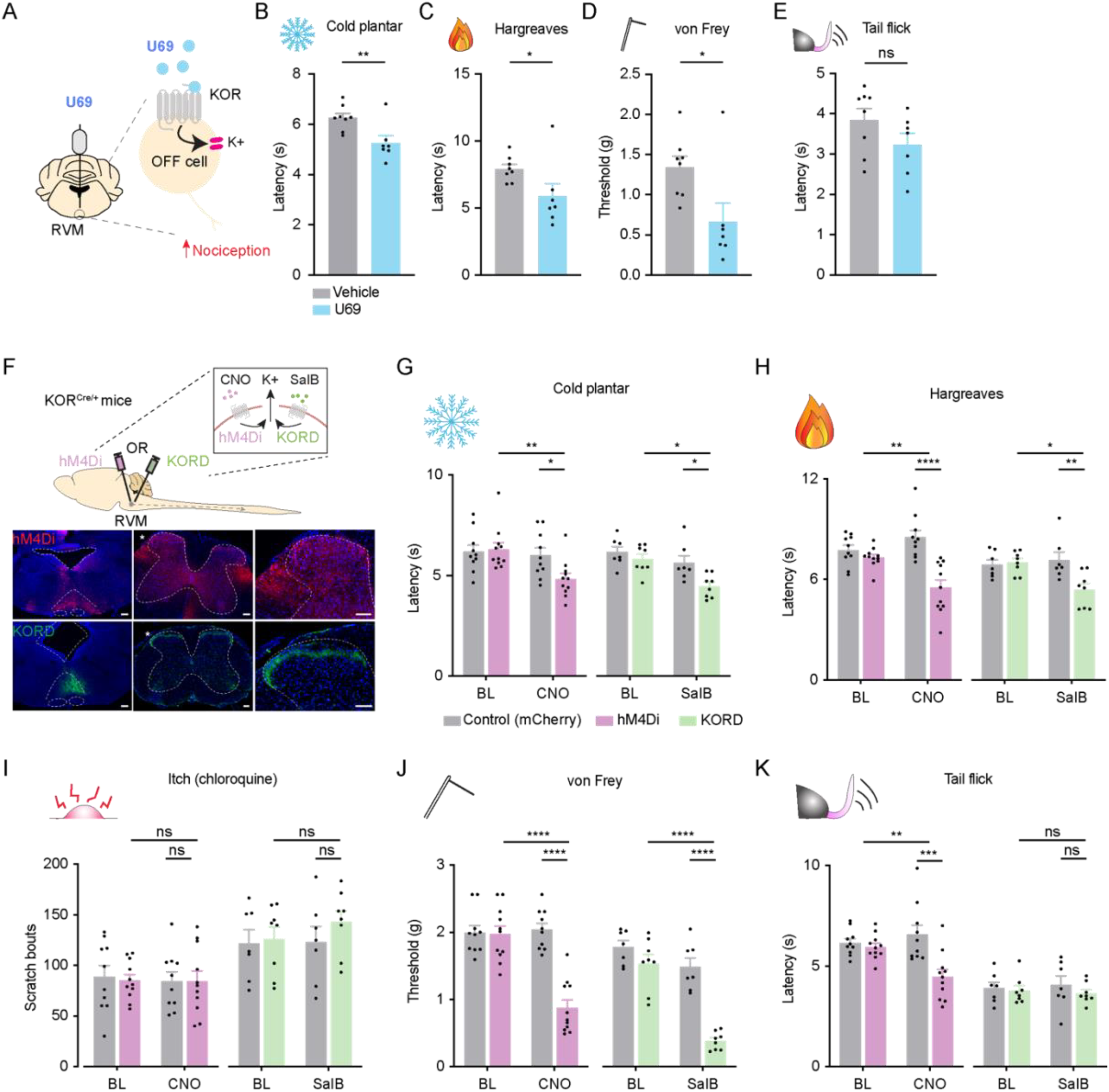
Pharmacological and chemogenetic inhibition of RVM^KOR^ neurons facilitates nociception. **(A)** Model to test whether the microinjection of a kappa agonist inhibits RVM^KOR^ neurons to facilitate nociception. **(B-E)** Effects of microinjection of U69,593 (40 ng) into the RVM on sensitivities to (B) cold plantar assay, **(C)** Hargreaves assay, **(D)** von Frey assay, and **(E)** tail-flick assay. Data are mean + SEM with dots representing individual mice (n = 8 vehicle, n = 7 U69,593 treated animals). Asterisks indicate the results of unpaired t-test (ns p>0.05, * p<0.05, ** p<0.01). **(F)** Model for how a chemogenetic strategy using hM4Di (pink) or KORD (green) DREADDs mimics the effects of pharmacological inhibition of RVM^KOR^ neurons. Representative images of the RVM and spinal cord following infection are shown. Scale bar = 50 μm. **(G-K)** Effect of chemogenetic inhibition of RVM^KOR^ neurons with CNO (IP 5 mg/kg) (pink, left) or salvinorin B (SC 10 mg/kg) (green, right) in mice receiving hM4Di or KORD, respectively, on **(G)** cold plantar assay, **(H)** Hargreaves, **(I)** chloroquine-evoked itch, **(J)** von Frey assay, and **(K)** tail-flick assay. Data are mean + SEM with dots representing individual mice (n = 10 hM4Di control, n = 11 hM4Di, n = 7 KORD control, n = 8 KORD mice). Asterisks indicate the results of two-way repeated measures ANOVA with Bonferroni’s correction (ns p>0.05, * p<0.05, ** p<0.01, *** p<0.001, **** p<0.0001).

Kappa agonists delivered to the RVM could affect either RVM neurons or presynaptic terminals from neurons that project into the RVM. Moreover, although kappa agonists traditionally reduce neuronal excitability through Giα signaling, they have also been shown to activate cellular kinases that facilitate neuronal activity.^41,42^ Thus, to more specifically address the effect of inhibiting RVM neurons that express KOR, we sought to chemogenetically inhibit RVM^KOR^ neurons using hM4Di and KORD DREADDs (Fig. 3F). We found that inhibition of RVM^KOR^ neurons with either hM4Di or KORD enhanced sensitivities to cold, thermal, and mechanical stimuli, but did not enhance chloroquine-evoked itch (Fig. 3G-J). In the tail-flick assay, only inhibition with hM4Di, but not KORD, reduced tail-flick latencies (Fig. 3K). Silencing of RVM^KOR^ neurons with hM4Di facilitated itch (Supplementary Fig. 4A), but did not exacerbate hyperalgesia following acute injury with intraplantar capsaicin (Supplementary Fig. 4B-E). Chemogenetic inhibition with either hM4Di or KORD had no effect on locomotor activity in an open field or rotarod assay, but KORD inhibition resulted in elevated responses to the acoustic startle test (Supplementary Fig. 4F-H). In agreement with our pharmacological inhibition of RVM^KOR^ neurons (Fig. 3B-D), these experiments suggest that RVM^KOR^ neurons are required for the descending inhibition of nociception, and their chemogenetic inhibition exposes a pro-nociceptive state.

### RVM^KOR^ neurons are required for stress-induced analgesia

In acute settings, stress inhibits pain^43^ and stress-induced analgesia has been blocked by the RVM microinjection of a KOR agonist.^6,44^ To test the contributions of RVM^KOR^ neurons to stress-induced analgesia, we used the forced swim model of stress-induced analgesia.^45,46^ We looked at stress-induced analgesia by using the hotplate assay and by examining nocifensive responses to intraplantar capsaicin (Fig. 4A). In control mice, forced swim decreased licking behavior and response latency on a hot plate, consistent with stress-induced analgesia. Upon activation RVM^KOR^ neurons with CNO alone (no stress), hot plate responses were decreased, similar to stress-induced analgesia, and stress had no further effect (Fig. 4B, C). Next, we examined the effect of inhibiting RVM^KOR^ neurons on stress-induced analgesia. Following inhibition with hM4Di, mice no longer exhibited stress-induced analgesia to intraplantar capsaicin or hotplate testing (Fig. 4D-F). Together, these results suggest that RVM^KOR^ neurons are necessary and sufficient for stress-induced analgesia.

**Figure 4.**
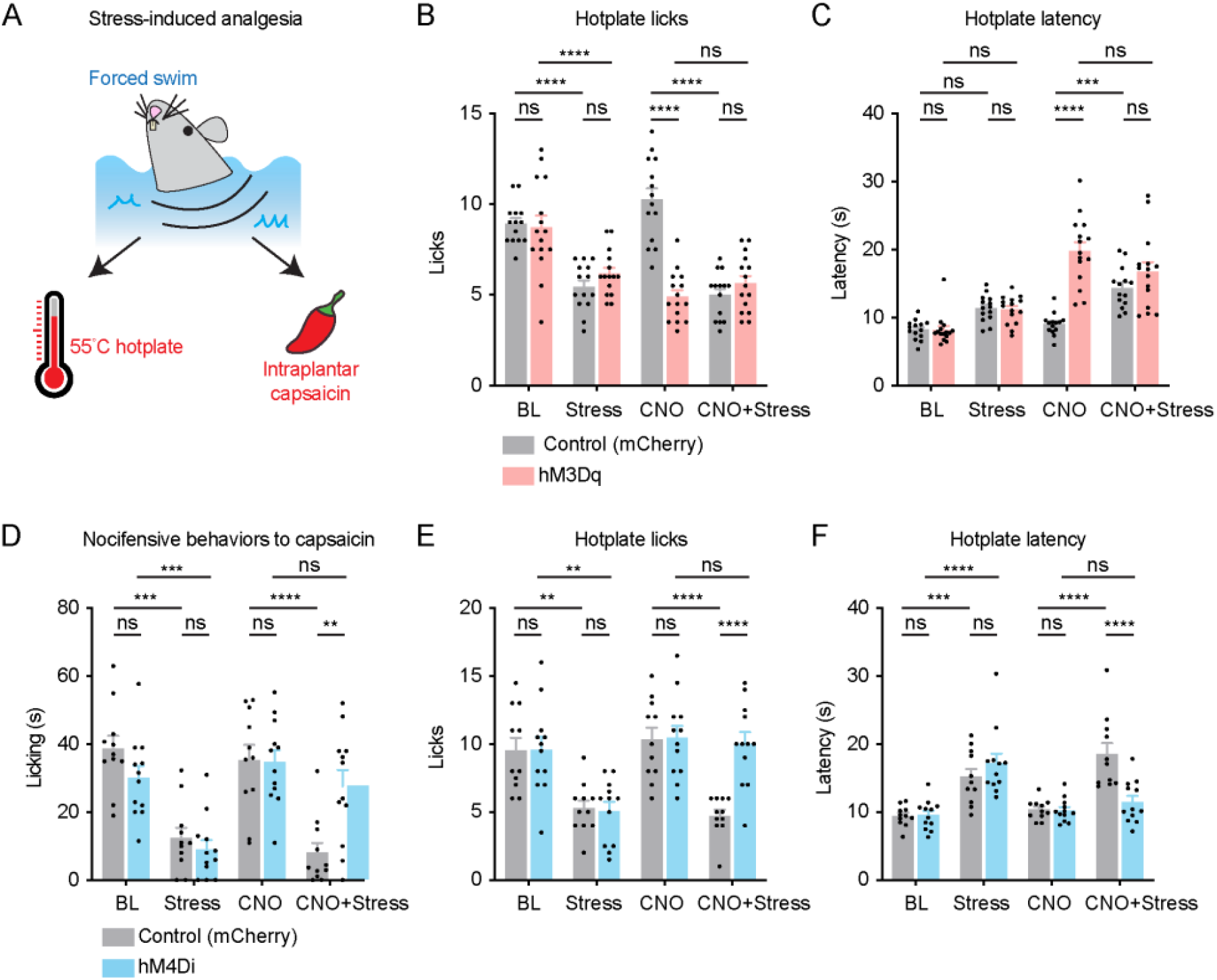
RVM^KOR^ neurons are required for stress-induced analgesia. **(A)** Model to test stress-induced analgesia. Mice were forced to swim for 3 minutes and later tested either on a hotplate (total licks and latency to lick or jump) or their licking responses to intraplantar capsaicin were quantified. **(B, C)** Effect of chemogenetic activation of RVM^KOR^ neurons (IP CNO 5 mg/kg) on responses to stress-induced analgesia as measured by **(B)** hotplate total licks and **(C)** hotplate licking latencies. Data are mean + SEM with dots representing individual mice (n = 14 control, n = 15 hM3Dq mice). Asterisks indicate the results of two-way repeated measures ANOVA with Bonferroni’s correction (ns p>0.05, *** p<0.001, **** p<0.0001). **(D-F)** Effect of chemogenetic inhibition of RVM^KOR^ neurons (IP CNO 5 mg/kg) on stress-induced analgesia as measured by **(D)** nocifensive responses to capsaicin, **(E)** hotplate total licks, and **(F)** hotplate licking latencies. Data are mean + SEM with dots representing individual mice (n = 11 control, n = 12 hM4Di mice). Asterisks indicate the results of two-way repeated measures ANOVA with Bonferroni’s correction (ns p>0.05, ** p<0.01, *** p<0.001, **** p<0.0001).

### Targeting of spinally-projecting RVM^KOR^ neurons

Given the robust innervation of the spinal cord by RVM^KOR^ neurons (Fig. 1B, Supplementary Fig. 2B, Fig. 2A, Supplementary Fig. 5A), and the role of the RVM in the descending modulation of pain, we sought to characterize the organization of RVM^KOR^ neurons that project to the spinal cord. Previous studies have reported variations in descending modulation between spinally-innervated lumbar segments, and uncovered the differential modulation of thermal sensitivity in the tail and foot.^47^ Furthermore, RVM inactivation has been shown to completely block allodynia in facial regions, but not plantar hind paw in an inflammatory pain model.^48^ Therefore, we examined the anatomical innervation of RVM^KOR^ neurons at different dermatomal segments. We used multi-color tracers with retrograde properties (AAVr-Ef1a-DIO-eYFP, AAVr-hSyn-DIO-mCherry, and CTB-647) and injected these tracers at different segmental levels to examine whether RVM^KOR^ neurons are somatotopically organized (Fig. 5A). We found that RVM^KOR^ neurons were frequently infected by both eYFP and mCherry, suggesting that the same RVM^KOR^ neurons innervate both the cervical and lumbar spinal segments. This observation suggests that these neurons are not somatotopically organized. From these experiments, we also determined that RVM^KOR^ neurons comprised 44.9 ± 1.5% of the total CTB-647-labeled spinally-projecting RVM neurons, which underscores that descending RVM^KOR^ neurons represent a significant fraction of total descending neurons from the RVM (Fig. 5A).

**Figure 5.**
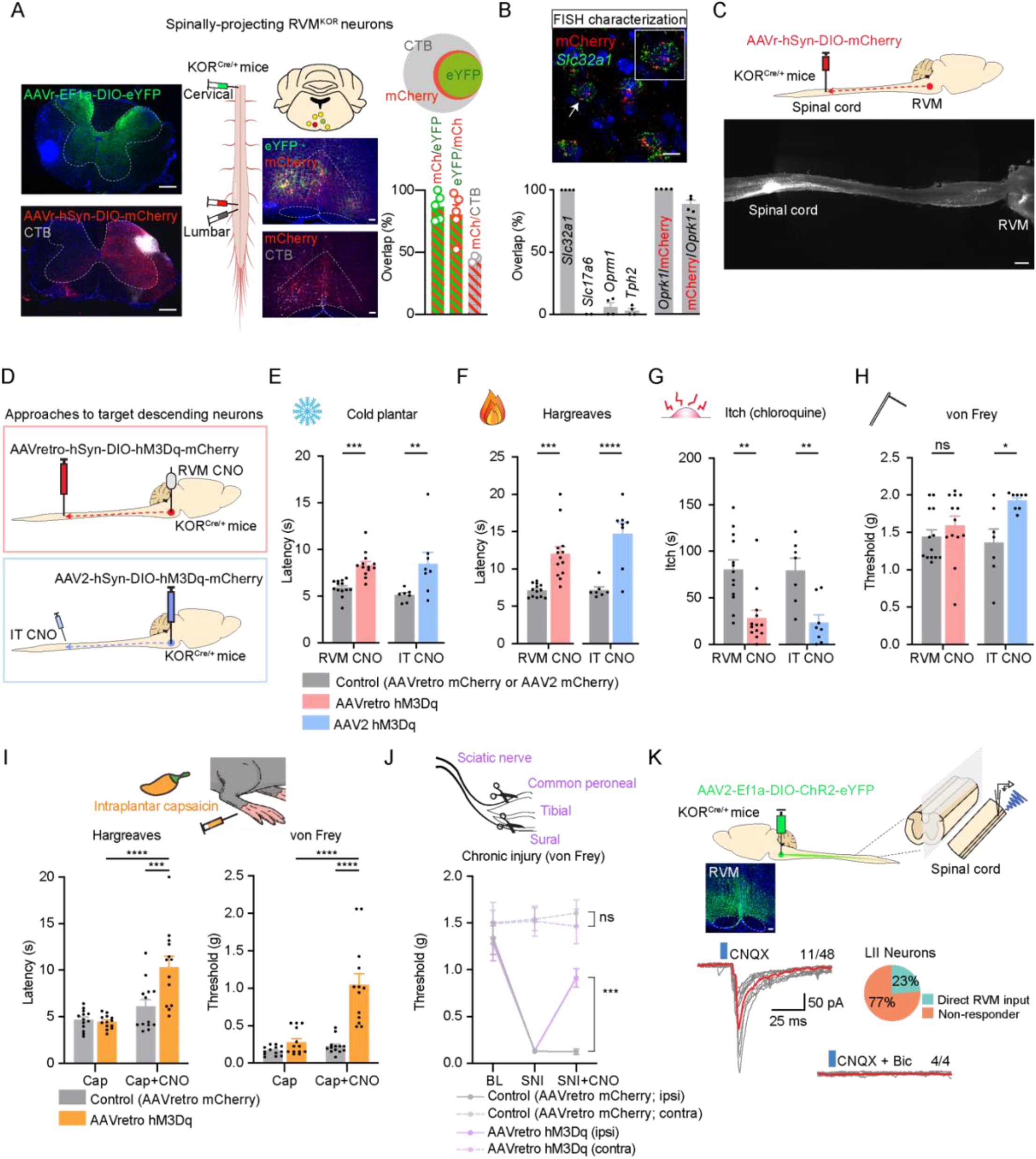
Activation of descending RVM^KOR^ neurons is analgesic. **(A)** Labeling of spinally-projecting RVM^KOR^ neurons using retrograde AAV and CTB introduced to the lumbar and cervical spinal cords. Data are mean + SEM with dots representing individual mice (n = 3-5). Scale bar = 50 μm. **(B)** FISH characterization of spinally-projecting RVM^KOR^ neurons. Data are mean + SEM with dots representing individual mice (n = 4). Scale bar = 50 μm. **(C)** Representative image of iDisco clearing of brainstem-spinal cord specimen following intraspinal injection of AAVr-FLEX-tdtomato. Scale bar = 1 mm **(D)** Approach to selectively activate spinally-projecting RVM^KOR^ neurons using two complementary approaches. One involves intraspinal AAVretro-hM3Dq and RVM CNO (1 mmol in 300 nL). The other involves injection of AAV2-hM3Dq into the RVM with the activation of descending input with intrathecal (IT) CNO (50 μg/kg). **(E to H)** Effect of chemogenetic activation of spinally-projecting RVM^KOR^ neurons using RVM CNO or IT CNO on responses to (E) cold plantar assay, (F) Hargreaves assay, (G) chloroquine-evoked itch, and (H) von Frey assay. Data are mean + SEM with dots representing individual mice (AAVretro: n = 13 controls, n = 13 hM3Dq; AAV2: n = 7 controls, n = 8 hM3Dq). Asterisks indicate the results of two-way repeated measures ANOVA with Bonferroni’s correction (ns p>0.05, * p<0.05, ** p<0.01, *** p<0.001, **** p<0.0001). **(I)** Effect of chemogenetic activation of spinally-projecting RVM^KOR^ neurons with RVM CNO on thermal and mechanical responses with intraplantar capsaicin-induced injury. Data are mean + SEM with dots representing individual mice (n = 13 controls, n = 13 hM3Dq). Asterisks indicate the results of two-way repeated measures ANOVA with Bonferroni’s correction (**** p<0.0001). **(J)** Effect of chemogenetic activation of spinally-projecting RVM^KOR^ neurons with RVM CNO on mechanical hypersensitivity with SNI. Data are mean ± SEM (n = 7 controls, n = 8 hM3Dq). Asterisks indicate the results of two-way repeated measures ANOVA with Bonferroni’s correction (ns p>0.05, *** p<0.001). **(K)** Electrophysiological recordings of lamina II spinal neurons to examine descending inputs from RVM^KOR^ neurons following the injection of AAV2-Ef1a-DIO-ChR2-eYFP in the RVM of KOR^Cre^ mice. Representative traces are shown. 11/48 cells in lamina II received direct input from RVM^KOR^ terminals following optogenetic activation (5 ms duration) and these responses are abolished in the presence of bicuculline (10 µM). All recordings were made in the presence of CNQX (10 µM).

Next, we molecularly characterized these spinally-projecting RVM^KOR^ neurons and found that they were virtually all GABAergic (based on their 100% overlap with *Slc32a1*) and that they exhibited limited overlap with other markers of RVM neurons (Fig. 5B). This finding is in striking contrast with RVM^KOR^ neurons as a heterogenous group (Fig. 2B), suggesting that spinally-projecting RVM^KOR^ neurons comprise a molecularly distinct population. Using this approach, we were also able to capture 88.1 ± 3.1% of *Oprk1* neurons in the RVM with 100% specificity (Fig. 5B). Furthermore, we performed whole-tissue clearing of intact brainstem-spinal cord specimens in which descending RVM^KOR^ neurons were labeled with intraspinal AAVr-FLEX-tdtomato to visualize this pathway (Fig. 5C, Supplementary Video 3). Using this technique, we were able to label the cell bodies of RVM^KOR^ neurons that project to the spinal cord. These anatomical and molecular approaches, therefore, allowed us to selectively target, characterize, and visualize descending RVM^KOR^ neurons.

### Descending RVM^KOR^ neurons inhibit itch and nociception through a spinal circuit

Although our experiments suggested that RVM^KOR^ neurons inhibit nociception and itch, the specific KOR neurons that were involved remained unclear. Our analysis had revealed that RVM^KOR^ neurons as a whole include GABAergic, glutamatergic and serotonergic subtypes; however, those that project to the spinal cord were exclusively GABAergic, and we hypothesized that it was these descending neurons that mediate the inhibition of nociception and itch. To test this idea, we used two approaches to manipulate spinally-projecting RVM^KOR^ neurons (Fig. 5D). In one set of experiments, we injected AAVr-hSyn-DIO-hM3Dq-mcherry into the lumbar spinal cord of KOR^Cre^ mice and implanted guide cannulas over the RVM, allowing us to infuse CNO directly into the RVM. In a complementary set of experiments, we tested animals receiving the diluted AAV2-hSyn-DIO-hM3Dq-mCherry into the RVM and administered intrathecal (IT) CNO, which allowed us to spatially restrict our chemogenetic activation to RVM^KOR^ spinal projections. We confirmed that chemogenetic activation, with either approach, did not result in unusual motor or arousal behaviors on an open field, rotarod, and acoustic startle test (Supplementary Fig. 3C-G). With both manipulations, activation of RVM^KOR^ cells inhibited responses to cold, thermal plantar testing, and chloroquine-evoked itch (Fig. 5E-G, Supplementary Fig. 3H-K), but only activation of descending fibers with IT CNO reduced mechanical responses to von Frey testing (Fig. 5H). Lastly, activation of these descending neurons inhibited capsaicin-induced thermal and mechanical hypersensitivities (Fig. 5I, Supplementary Fig. 3L, M) as well as mechanical allodynia following chronic injury with spared-nerve injury (Fig. 5J). These findings highlight the key role of spinally-projecting RVM^KOR^ neurons in the inhibition of nociception, itch, and acute and chronic pain.

To determine the nature of RVM^KOR^ inputs to the spinal cord, we conducted recordings in spinal cord slices of KOR^Cre^ animals that had received AAV2-hSyn-DIO-ChR2-eYFP in the RVM (Fig. 5K, Supplementary Fig. 4B-E). We patched random cells in lamina II and found that nearly a quarter of these neurons received direct inhibitory inputs from RVM^KOR^ neurons upon optogenetic activation (Fig. 5K). We further validated that the nature of this input was inhibitory by blockade with bicuculline (Fig. 5K). Additionally, the latency and jitter of optically evoked IPSCs (Supplementary Fig. 4B, D) was consistent with the idea that inhibitory input from RVM^KOR^ neurons onto lamina II neurons is monosynaptic. Together, these findings suggest that the anti-nociceptive effects of activating RVM^KOR^ neurons may occur through direct inhibition of spinal neurons.

We then applied our intersectional viral and pharmacological approach to determine the effects of inhibiting descending RVM^KOR^ neurons. We selectively removed the kappa-opioid receptor from descending neurons by injecting AAVr-hSyn-eGFP-Cre into the lumbar spinal cord of KOR^fl/fl^ mice (Fig. 6A). Consistent with our previous findings that microinjection of U69,593 into the RVM is pro-nociceptive in wild type mice (Fig. 3B-D), administration of this kappa agonist also generally unveiled a pro-nociceptive state upon cold and mechanical testing, but did not result in thermal hypersensitivity (Fig. 6B-E). As expected (Fig. 3E), local injection of U69,593 did not affect the tail flick test (Fig. 6E). Following the viral-mediated removal of the kappa-opioid receptor from descending RVM neurons, we found that kappa agonism no longer elicited cold and mechanical hypersensitivity (Fig. 6B, D), highlighting that expression of the kappa-opioid receptor on descending RVM^KOR^ neurons, and not non-spinally-projecting RVM^KOR^ neurons or presynaptic terminals from neurons that project into the RVM, is required for the pro-nociceptive effects following kappa agonism.

**Figure 6.**
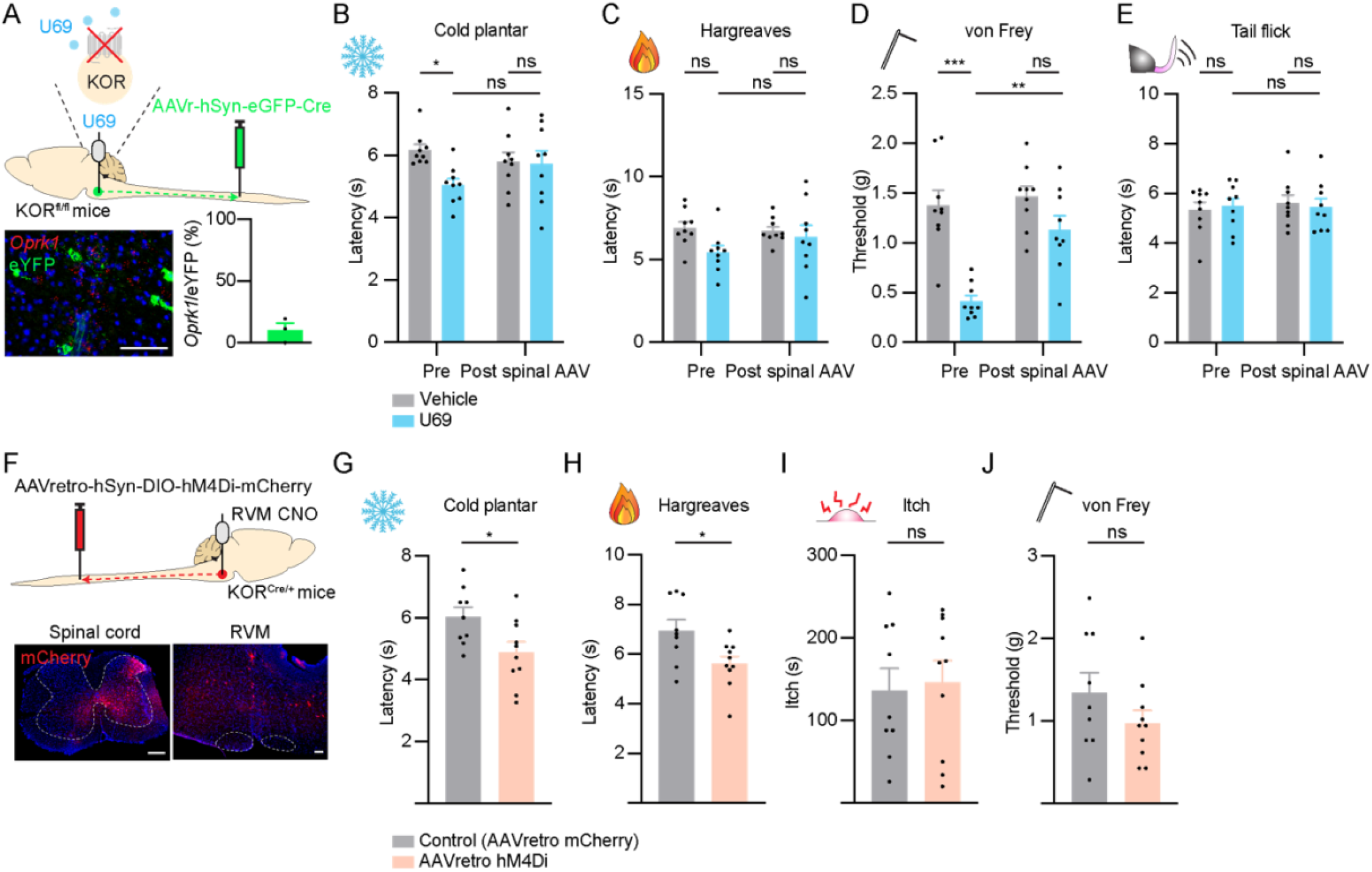
Inhibition of descending RVM^KOR^ neurons facilitates nociception. **(A)** Approach to selectively remove the kappa-opioid receptor from descending RVM neurons. KOR^fl/fl^ animals received AAVr-hSyn-eGFP-Cre into the lumbar spinal cord. Validation of the deletion of KOR within the RVM was determined by FISH. Data are mean + SEM with dots representing individual mice (n = 3 mice). Scale bar = 50 μm. **(B-E)** Effects of microinjection of U69,593 (40 ng) into the RVM before and after Cre-mediated deletion of KOR within the RVM. Sensitivities to **(B)** cold plantar assay, **(C)** Hargreaves assay, **(D)** von Frey assay, and **(E)** tail-flick assay were tested. Data are mean + SEM with dots representing individual mice (n = 9 vehicle, n = 7 U69,593 treated animals). Asterisks indicate the results of two-way ANOVA with Bonferroni’s correction (ns p>0.05, * p<0.05, ** p<0.01, *** p<0.001). **(F)** Strategy to selectively inhibit descending RVM^KOR^ neurons using AAVr-hSyn-DIO-hM4Di-mCherry into the lumbar spinal cord followed by implantation of a guide cannula over the RVM to permit the local infusion of CNO. Representative targeting of the spinal cord and labeling of cell bodies within the RVM are shown. Scale bar = 50 μm. **(G-J)** Effect of chemogenetic inhibition of RVM^KOR^ neurons with local CNO (1 mmol in 300 nL) on **(G)** cold plantar assay, **(H)** Hargreaves, **(I)** chloroquine-evoked itch, and **(J)** von Frey assay. Data are mean + SEM with dots representing individual mice (n = 9 AAVretro control, n = 10 AAVretro hM4Di mice). Asterisks indicate the results of unpaired t-test (ns p>0.05, * p<0.05).

In a complementary set of experiments, we selectively inhibited descending RVM^KOR^ neurons by viral delivery of AAVretro-hSyn-DIO-hM4Di-mCherry into the lumbar spinal cord of KOR^Cre^ mice and local CNO into the RVM (Fig. 6F). The chemogenetic inhibition of descending RVM^KOR^ neurons increased sensitivity to cold and Hargreaves testing but did not affect itch or mechanical withdrawal (Fig. 6G-J). Furthermore, selective inhibition of spinally-projecting RVM^KOR^ neurons attenuated stress-induced analgesia to capsaicin and hotplate testing (Supplementary Fig. 4I-K).

Together, our combinatorial pharmacological and viral approach to selectively inhibit descending RVM^KOR^ neurons reinforces the idea that these neurons function to inhibit nociception.

### Dynorphin signaling between the PAG and RVM bidirectionally modulates nociception

Next, we investigated potential inputs to RVM^KOR^ neurons. As dynorphin is the endogenous peptide for the kappa-opioid receptor, we injected Cre-dependent retrograde viral tracers into the RVM in Dyn^Cre^ mice to identify areas that could provide dynorphinergic inputs to the RVM (Fig. 7A). Robust labeling was identified in many areas, including the parabrachial nucleus (PBN), dorsal reticular nucleus (Drt), and the agranular insular area (AIp); however, the most robust labeling was in the ventrolateral periaqueductal gray (vlPAG) (Fig. 7A). Previous studies have highlighted the RVM as a necessary component within the descending modulatory circuit activated by stimulation of the PAG. For example, lesioning or inactivation of the RVM (with local anesthetics) blocks the anti-nociceptive action of PAG stimulation.^49–51^ Therefore, we focused on RVM-projecting PAG^Dyn^ neurons as possible inputs to RVM^KOR^ cells. We characterized RVM-projecting PAG^Dyn^ neurons using CTB and AAVr-DIO-tdt in Dyn^Cre^ mice (Fig. 7B) and found that PAG^Dyn^ neurons represent 10.4 ± 4.0% of PAG neurons that project to the RVM. Using a complementary approach involving retrograde viral labeling and FISH, we found that this RVM-projecting population represented 32.3 ± 2.1% of PAG^Dyn^ neurons. Additionally, glutamatergic (*Slc17a6-*expressing) neurons comprised 99.3 ± 0.7% of all RVM-projecting PAG^Dyn^ neurons (Fig. 7C). Lastly, PAG^Dyn^ neurons represented a population that was molecularly distinct from PAG^Tac1^ and PAG^Th^ neurons that have recently been described in the modulation of itch and nociceptive behaviors, respectively^52,53^ (Supplementary Fig. 6A-C). Together, these findings identify a unique population of glutamatergic PAG^Dyn^ neurons that project to the RVM.

**Figure 7.**
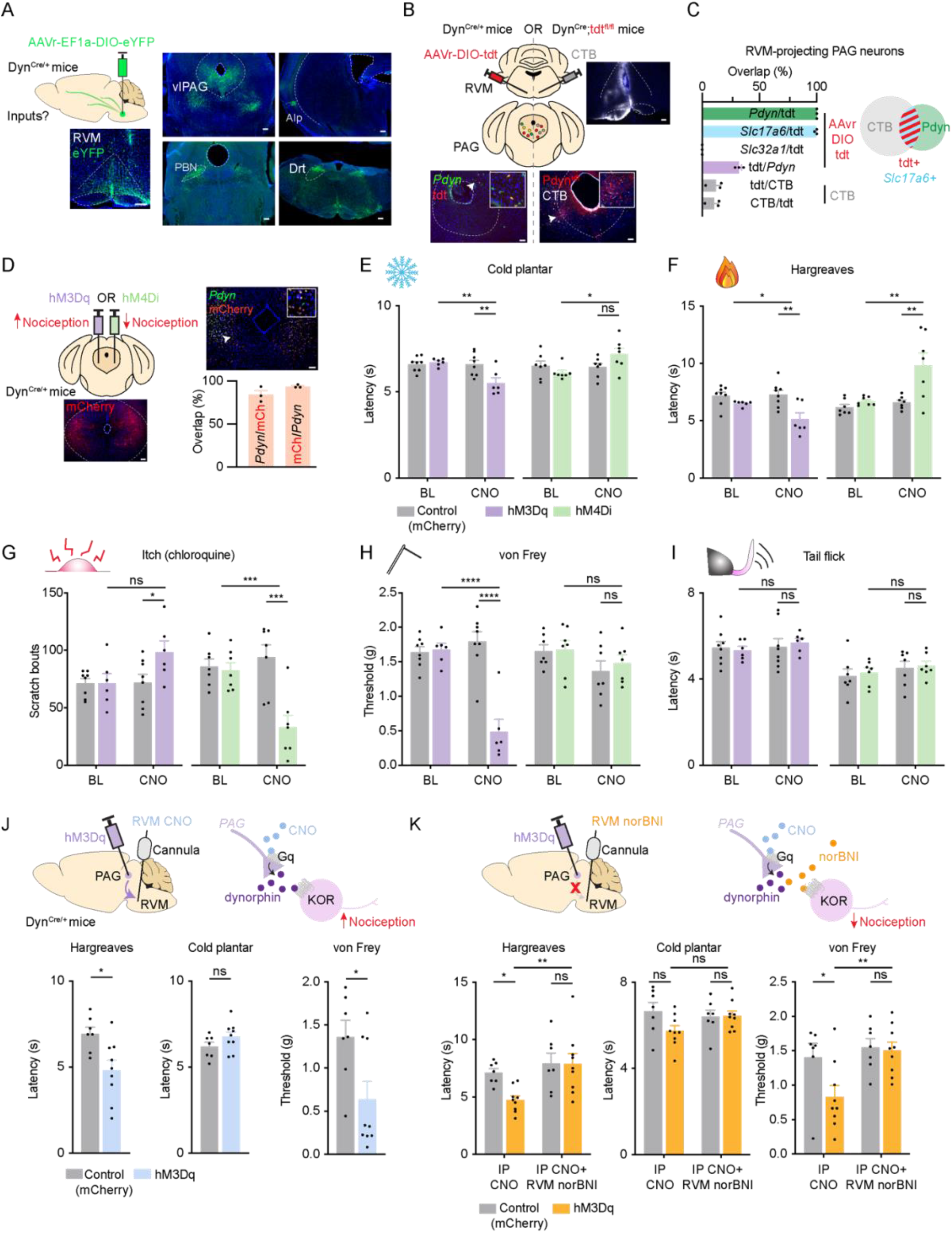
PAG^Dyn^ neurons provide input to the RVM and bidirectionally modulate pain. **(A)** Experimental strategy to trace dynorphinergic inputs to the RVM. Robust labeling was observed in the PAG of Dyn^Cre^ mice receiving AAVr-Ef1a-DIO-eYFP in the RVM. vlPAG:ventrolateral periaqueductal gray, PBN: parabrachial nucleus, AIp: agranular insular area, Drt: dorsal reticular nucleus. Scale bar = 50 μm. **(B)** Approach to characterize dynorphin neurons projecting to the RVM. In one experiment, Dyn^Cre^ mice received Cre-dependent AAVr-DIO-tdt in the RVM and RVM-projecting neurons were further characterized using FISH. In a separate experiment, Dyn^Cre^;tdtomato mice received CTB and back labeled CTB neurons were quantified. Scale bar = 50 μm. **(C)** Summary of RVM-projecting PAG^Dyn^ neurons characterization. Data are mean + SEM with dots representing individual mice (n = 3). **(D)** Experimental strategy and effect of bidirectional modulation of PAG^Dyn^ neurons using chemogenetic activation (purple) and inhibition (green) of with IP CNO (5 mg/kg). Validation of viral targeting using FISH is shown. Data are mean + SEM with dots representing individual mice (n = 3 mice). Scale bar = 50 μm. **(E-I**) Effect of chemogenetic activation with hM3Dq (purple, left) and inhibition with hM4Di (green, right) on **(E)** cold plantar assay, **(F)** Hargreaves assay, **(G)** chloroquine-evoked itch, **(H)** von Frey assay, and **(I)** tail-flick assay. Data are mean + SEM with dots representing individual mice (hM3Dq: n = 8 control, n = 6 hM3Dq; hM4Di: n = 7 control, n = 7 hM4Di mice). Asterisks indicate the results of two-way repeated measures ANOVA with Bonferroni’s correction (ns p>0.05, * p<0.05, ** p<0.01, *** p<0.001, **** p<0.0001). **(J)** Experimental approach to selectively activate PAG^Dyn^ projections to the RVM by delivery of RVM CNO (1 mmol in 300 nL). Effect of chemogenetic activation on responses to Hargreaves, cold plantar, and von Frey testing. Data are mean + SEM with dots representing individual mice (n = 7 control, n = 9 hM3Dq mice). Asterisks indicate the results of unpaired t-test (ns p>0.05, * p<0.05). **(K)** Experimental approach to block dynorphin input from chemogenetically-activated PAG^Dyn^ neurons by delivery of RVM norBNI (100 ng in 250 nL) and its effects on Hargreaves, cold plantar, and von Frey testing. Data are mean + SEM with dots representing individual mice (n = 7 control, n = 9 hM3Dq mice). Asterisks indicate the results of two-way repeated measures ANOVA with Bonferroni’s correction (ns p>0.05, * p<0.05, ** p<0.01, *** p<0.001, **** p<0.0001).

Recent work has revealed that glutamatergic and GABAergic PAG neurons divergently modulate nociception and itch.^54,55^ We bidirectionally modulated the dynorphin population of PAG neurons using Cre-dependent hM3Dq and hM4di in Dyn^Cre^ mice to activate and inhibit these neurons, respectively (Fig. 7D). Chemogenetic activation and inhibition with IP CNO bidirectionally modulated thresholds to cold plantar and Hargreaves testing (Fig. 7E, F) and scratching behaviors in response to chloroquine (Fig. 7G). Although activation of PAG^Dyn^ neurons reduced mechanical thresholds, inhibition did not elevate thresholds above baseline (Fig. 7H). Neither activation nor inhibition of PAG^Dyn^ neurons affected latencies to tail-flick (Fig. 7I). Overall, we found that activation of PAG^Dyn^ neurons generally facilitated nociception and itch, whereas inhibition of these neurons reduced sensitivities to cold, thermal, and itch responses.

We also confirmed that PAG^Dyn^ neurons facilitated nociception through their input to the RVM by implanting cannulas in the RVM for the local delivery of CNO (Fig. 7J). This allowed us to selectively activate RVM-projecting PAG^Dyn^ neurons. We found that the selective chemogenetic activation of this pathway generally recapitulated our findings following IP CNO (Fig. 7F, H) and facilitated sensitivity to thermal and mechanical testing, but not to cold testing (Fig. 7J).

To test more directly whether PAG^Dyn^ neurons could exert their effects through kappa signaling in the RVM, we used a pharmacological approach to block dynorphin input to the RVM from PAG^Dyn^ neurons (Fig. 7K). We found that IP CNO reduced latencies to Hargreaves testing and mechanical thresholds in the von Frey assay. However, microinjection of norBNI, a kappa antagonist, prior to the administration of IP CNO blocked the development of thermal and mechanical hypersensitivity (Fig. 7K). Thus, these data suggest that PAG^Dyn^ neurons facilitate nociception through kappa signaling in the RVM.

### Mu agonists disinhibit RVM^KOR^ neurons

Previous models have suggested that morphine inhibits nociception, at least in part, through the indirect activation of OFF cells in the RVM, potentially through a mechanism of disinhibition. In particular, OFF cells are proposed to be directly inhibited by kappa agonists, but disinhibited by mu agonists.^23–25^ We set out to test whether the morphine-active cells are *Oprk1*-expressing neurons. As expected, microinjection of morphine into the RVM increased response latencies to thermal stimulation in the Hargreaves assay (Fig. 8A). This anti-nociception was accompanied by the induction of *Fos* in 40.2 ± 9.9% of *Oprk1*-expressing neurons, which represented 40.9 ± 8.3% of *Fos*-expressing cells (Fig. 8B). When we recorded from spinally-projecting RVM^KOR^ neurons by injecting AAVr-FLEX-tdtomato into the spinal cord of KOR^Cre^ mice, we found that DAMGO substantially reduced the frequency of miniature inhibitory postsynaptic currents (mIPSCs) in a subset of these cells (Fig. 8C, Supplementary Fig. 5F, G). Importantly, no RVM^KOR^ neurons showed a direct effect of DAMGO compared to baseline, suggesting that they are not ON cells (Supplementary Fig. 5G). Thus, our findings support the idea that morphine enhances the activity of RVM^KOR^ neurons to produce anti-nociception, implicating RVM^KOR^ neurons as OFF cells (Fig. 8D).

**Figure 8.**
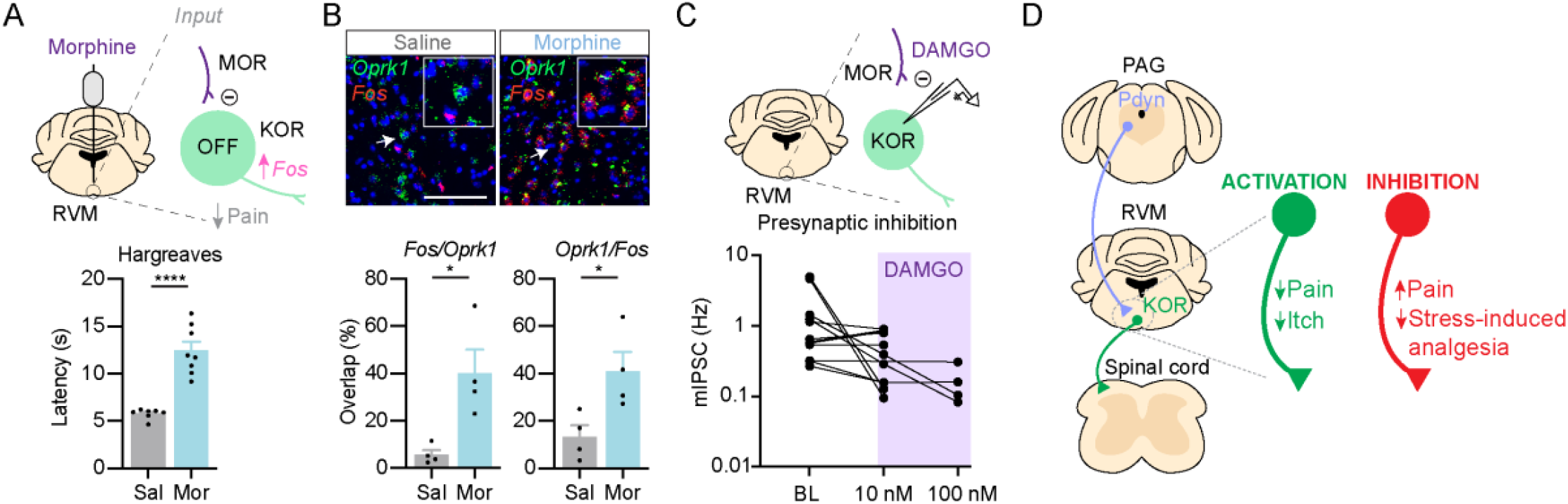
Model for RVM^KOR^ neurons as OFF cells. **(A)** Model to test whether RVM^KOR^ neurons are OFF cells by examining if microinjection of morphine disinhibits RVM^KOR^ neurons to produce analgesia. Effect of microinjection of morphine (1 μg in 250 nL) into the RVM on latencies in the Hargreaves assay. Data are mean + SEM with dots representing individual mice (n = 7 saline, n = 8 morphine treated animals). Asterisks indicate the results of unpaired t-test (**** p<0.0001). **(B)** *Fos* expression is enriched in *Oprk1* neurons following microinjection of morphine into the RVM. Data are mean + SEM with dots representing individual mice (n = 4 saline, n = 4 morphine treated animals). Scale bar = 50 μm. Asterisks indicate the results of unpaired t-test (* p<0.05). **(C)** Recordings of labeled spinally-projecting RVM^KOR^ neurons revealed a reduction in inhibitory synaptic transmission in the presence of DAMGO (10 and 100 nM). Out of 11 cells, 6 showed a reduction in mIPSC frequency but only 4 of those were significant at p < 0.05 by a Kruskal-Wallis test. **(D)** Model for the role of RVM^KOR^ neurons in the descending modulation of nociception. PAG^Dyn^ neurons provide inputs to the RVM. RVM^KOR^ neurons target the spinal cord and bidirectionally modulate nociception.

## Discussion

Our study identifies the role of RVM^KOR^ neurons in the inhibition of pain and itch. Previous work has also implicated RVM^KOR^ neurons as OFF cells.^23–25^ However, there was controversy as to whether RVM neurons responsive to kappa agonists could be pro-or anti-nociceptive.^31,32^ One limitation of previous studies is that pharmacological manipulation cannot distinguish between direct effects on RVM neurons^23–25^ or presynaptic terminals.^31,32^ Our neurochemical characterization of RVM^KOR^ neurons also reveals that they comprise a heterogeneous population; yet our ability to capture and manipulate spinally-projecting RVM^KOR^ neurons identifies them as a molecularly distinct subgroup of all RVM^KOR^ neurons. We found that the targeting of a small number of RVM^KOR^ neurons that project to the spinal cord was sufficient to produce anti-nociception and revealed that RVM^KOR^ neurons are required for stress-induced analgesia.

We found that all RVM^KOR^ neurons that project to the spinal cord are GABAergic. However, the KOR-expressing subset only represents 40% of all spinally-projecting GABAergic neurons in the RVM, suggesting that spinally-projecting GABAergic neurons likely represent numerous cell types. Consistent with this idea, various groups that have modulated GABAergic RVM neurons had different findings. One study, using very similar approaches to those used in ours, uncovered the role of RVM^Vgat^ neurons in the facilitation of mechanical, but not heat, nociception^19^; however, another study found RVM^Gad2^ neurons (which capture a subpopulation of Vgat neurons) to be involved in the inhibition of heat nociception.^18^ Our manipulation of RVM^Gad2^ neurons is consistent with the latter findings, where we found that activation of these cells was also anti-nociceptive. Thus, it is likely that GABAergic neurons in the RVM span several functional subgroups and that differences in behavioral findings may reflect the relative subpopulations that are captured with the viral approach used. Future efforts involving intersectional genetic approaches will be required to identify and directly manipulate unique RVM subpopulations in freely behaving mice.

Several groups have examined the contribution of the RVM to the state-dependence of somatosensation, including nociception.^11,56–60^ Our findings suggest that RVM^KOR^ neurons are engaged during states of acute stress and that they mediate stress-induced analgesia. Such a protective mechanism may explain, for example, how in emergency situations, the survival of an organism depends on its ability to appropriately respond to nociceptive stimuli. Unexpectedly, our experiments also uncovered the possible role of other KOR neurons in the brainstem that could drive profound and diverse functions in movement and arousal. The precise identities and roles remain to be further elucidated.

Dynorphin signaling within the brain has been shown to be involved in many behaviors, including stress, addiction, and analgesia.^61,62^ We found that the selective activation and inhibition of dynorphin neurons in the PAG facilitates and inhibits nociception, respectively. Using pharmacological approaches, we determined that these PAG neurons influence nociception through their release of dynorphin within the RVM. This finding, together with the observation that we and others have made that RVM neurons are sensitive to kappa agonism and antagonism, suggests that the descending modulation of pain is influenced by dynorphin signaling between the PAG and RVM. Identification of this dynorphinergic pathway may provide opportunities to take advantage of the body’s endogenous opioid system to manage pain. Harnessing the endogenous pain-modulatory system may reduce the need for pharmacological treatments, such as opioids, with fewer side effects and improved outcomes for the treatment of pain disorders.

## Competing interests

The authors report no competing interests.

**Supplementary Figure 1.**
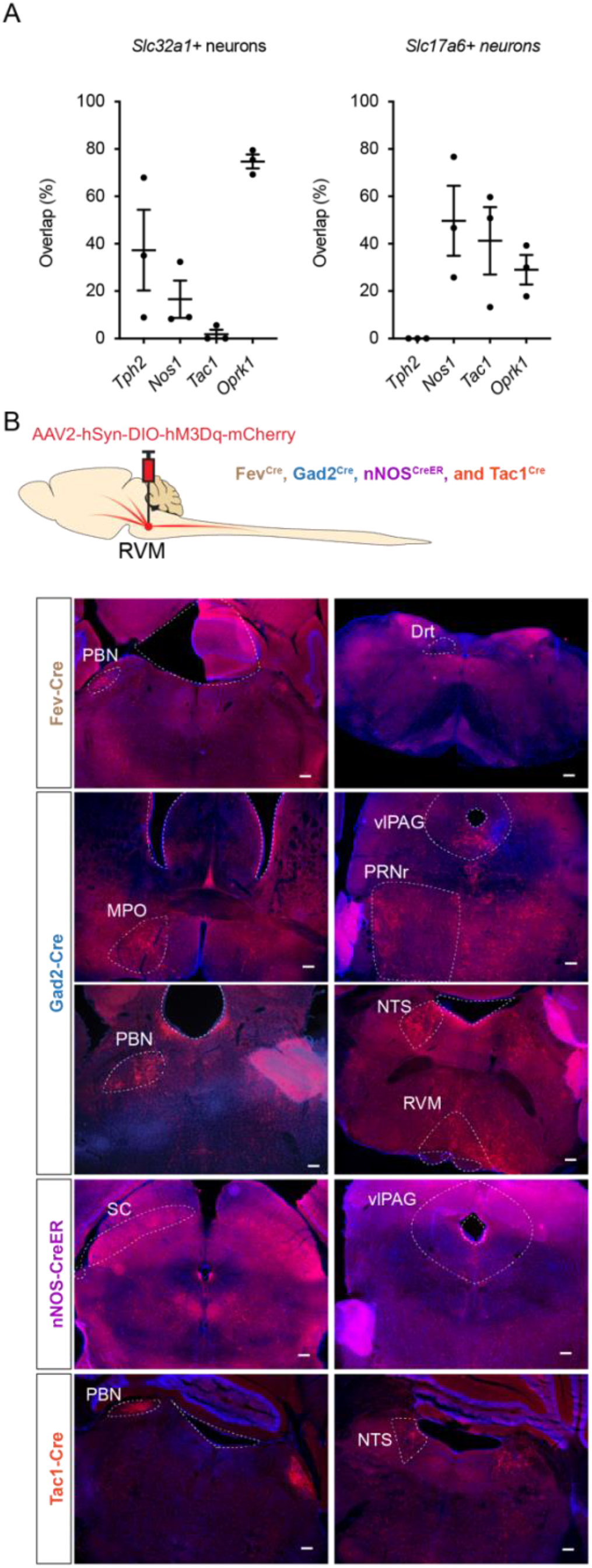
Tracing of RVM cell types to supraspinal structures. **(A)** FISH characterization of *Tph2, Nos1, Tac1* and *Oprk1* RVM neurons and their overlap with inhibitory (*Slc32a1)* and excitatory (*Slc17a6)* markers. *The same data pertaining to the characterization of KOR neurons in glutamatergic and GABAergic neurons are also shown in Figure 2B*. **(B)** Experimental approach to deliver Cre-dependent hM3Dq-mCherry to visualize labeled RVM neurons and their supraspinal projections across Fev^Cre^, Gad2^Cre^, nNos^CreER^, and Tac1^Cre^ alleles. Representative supraspinal targets are shown. PBN: parabrachial nucleus, Drt: dorsal reticular nucleus, MPO: medial preoptic area, vlPAG: ventrolateral periaqueductal gray, PRNr: pontine reticular nucleus, NTS: nucleus solitary tract, RVM: rostral ventromedial medulla, and SC: superior colliculus. Scale bar = 50 μm.

**Supplementary Figure 2.**
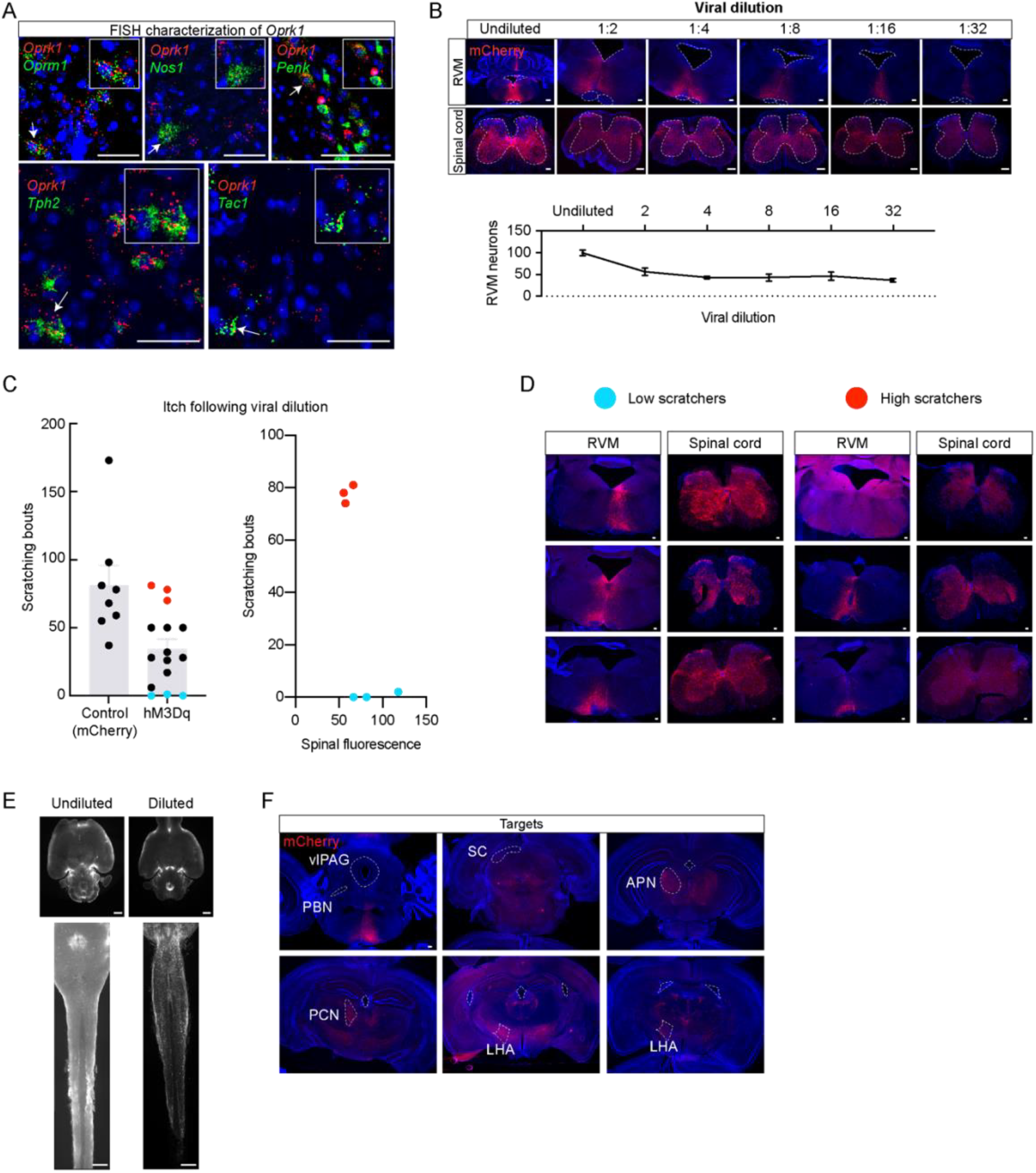
Restricted targeting of RVM^KOR^ neurons reveals their contribution to somatosensation. **(A)** Representative images of FISH characterization of RVM^KOR^ neurons for the quantifications shown in Figure 2B. Scale bar = 50 μm. **(B)** Serial dilutions of the hM3Dq-mCherry virus (original titer: 8.6×10^10^) captures smaller populations of RVM^KOR^ neurons. Scale bar = 50 μm. **(C)** A separation in the distribution of animals receiving 32-fold diluted hM3Dq reveals animals that behave comparably to controls (red) and animals with abolished itch responses to chloroquine (light blue). On the right, the mean density of fluorescence in the corresponding spinal cord tissues for the labeled animals is correlated to their scratching behaviors. **(D)** Representative RVM and spinal cord images of animals coded in (C). Scale bar = 50 μm. **(E)** iDisco cleared brain and brainstem-spinal samples in animals receiving viral injections into the RVM with and without dilution. Scale bar = 1 mm **(F)** Representative coronal sections of downstream targets of RVM^KOR^ neurons in animals receiving dilute titers of virus are shown (compare to Figure 1G). Scale bar = 50 μm.

**Supplementary Figure 3.**
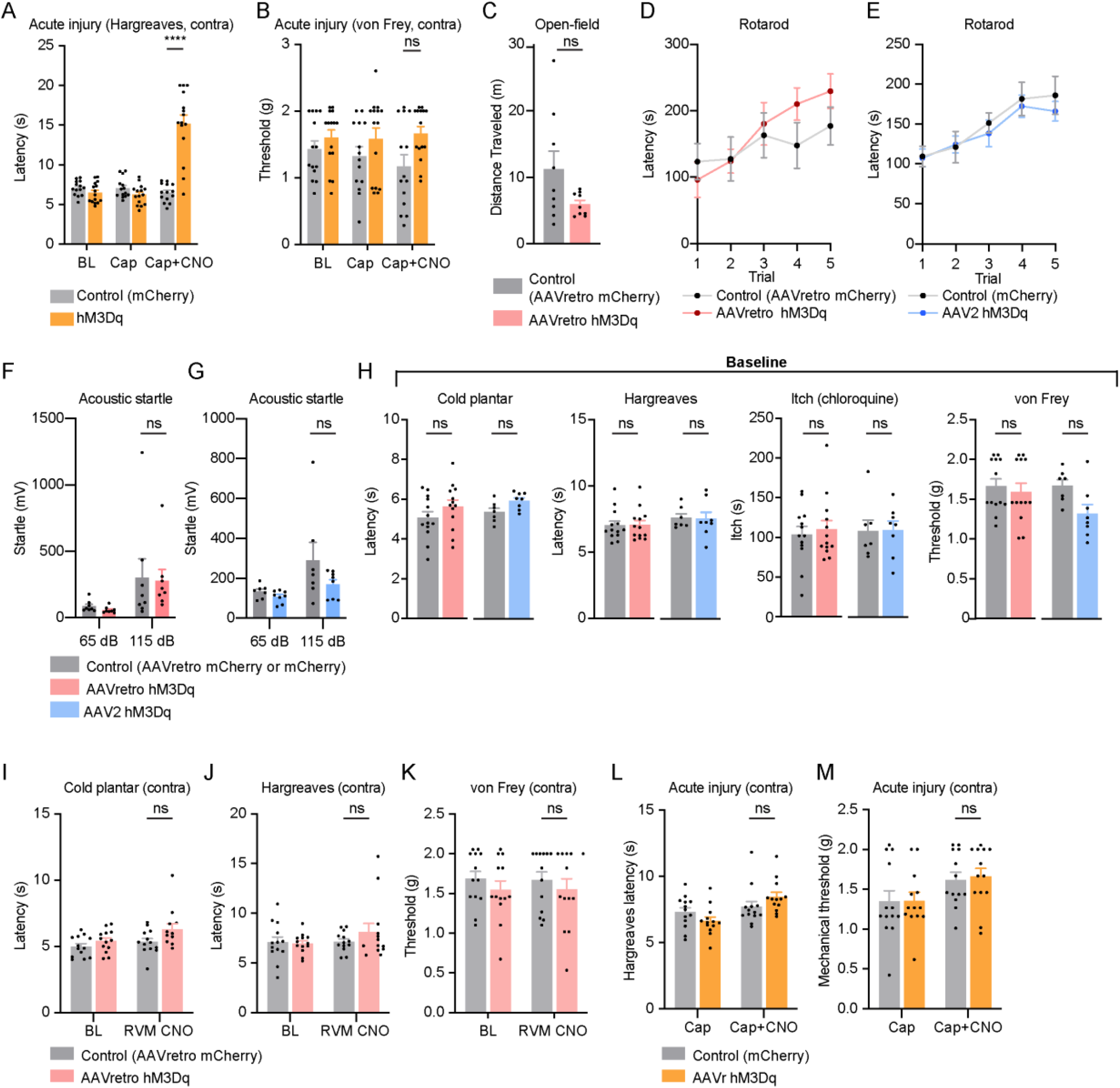
Supplementary chemogenetic activation behavioral data. **(A, B)** Effect of chemogenetic activation of RVM^KOR^ on (A) thermal and (B) mechanical responses following intraplantar capsaicin-induced injury on the contralateral, uninjured paw. Data are mean + SEM with dots representing individual mice (n = 14 controls, n = 15 hM3Dq). Asterisks indicate the results of two-way repeated measures ANOVA with Bonferroni’s correction (* p<0.05, **** p<0.0001). **(C)** Activation of descending RVM^KOR^ neurons with local CNO does not affect locomotor activity in an open field test. Data are mean + SEM with dots representing individual mice (n = 9 control, n = 9 AAVretro hM3Dq mice). NS indicates the results of one-way ANOVA with Bonferroni’s correction (ns p>0.05). **(D-G)** Activation of descending RVM^KOR^ neurons with either local CNO (D,F) or intrathecal CNO **(E**,**G)** did not affect performance on a rotarod or responses to acoustic startle. Data are mean ± SEM (D-E) and mean + SEM (F-G) (n = 9 AAVretro control, n = 9 AAVretro hM3Dq; n = 7 control hM3Dq, n = 8 hM3Dq mice. NS indicates the results of two-way ANOVA with Bonferroni’s correction (ns p>0.05). **(H)** Baseline responses to nociception and itch behaviors in animals receiving intraspinal AAVr-hSyn-DIO-hM3Dq-mCherry or intra-RVM AAV2-hSyn-DIO-hM3Dq-mCherry and their controls (AAVr-hSyn-DIO-mCherry or AAV2-hSyn-DIO-mCherry, respectively). Data are mean + SEM with dots representing individual mice (AAVretro: n = 13 controls, n = 13 hM3Dq; AAV2: n = 7 controls, n = 8 hM3Dq). Asterisks indicate the results of two-way repeated measures ANOVA with Bonferroni’s correction (ns p>0.05). *Baseline data for AAV2-hM3Dq are identical to those also shown in main Figure 2D-G, and are provided again here for comparison to the AAVretro-hM3Dq baselines*. **(I-K)** Effect of chemogenetic activation of spinally-projecting RVM^KOR^ neurons on the paw contralateral to the intraspinal viral injection and contralateral to the injury measured by **(I)** cold plantar assay, **(J)** Hargreaves assay, and **(K)** von Frey assay. Data are mean + SEM with dots representing individual mice (n = 13 controls, n = 13 hM3Dq). NS indicates the results of two-way repeated measures ANOVA with Bonferroni’s correction (ns p>0.05). **(L, M)** Effect of chemogenetic activation of spinally-projecting RVM^KOR^ neurons on the paw contralateral to the intraspinal viral injection and contralateral to the injury measured by **(L)** thermal and **(M)** mechanical testing. Data are mean + SEM with dots representing individual mice (n = 13 controls, n = 13 hM3Dq). NS indicates the results of two-way repeated measures ANOVA with Bonferroni’s correction (ns p>0.05).

**Supplementary Figure 4.**
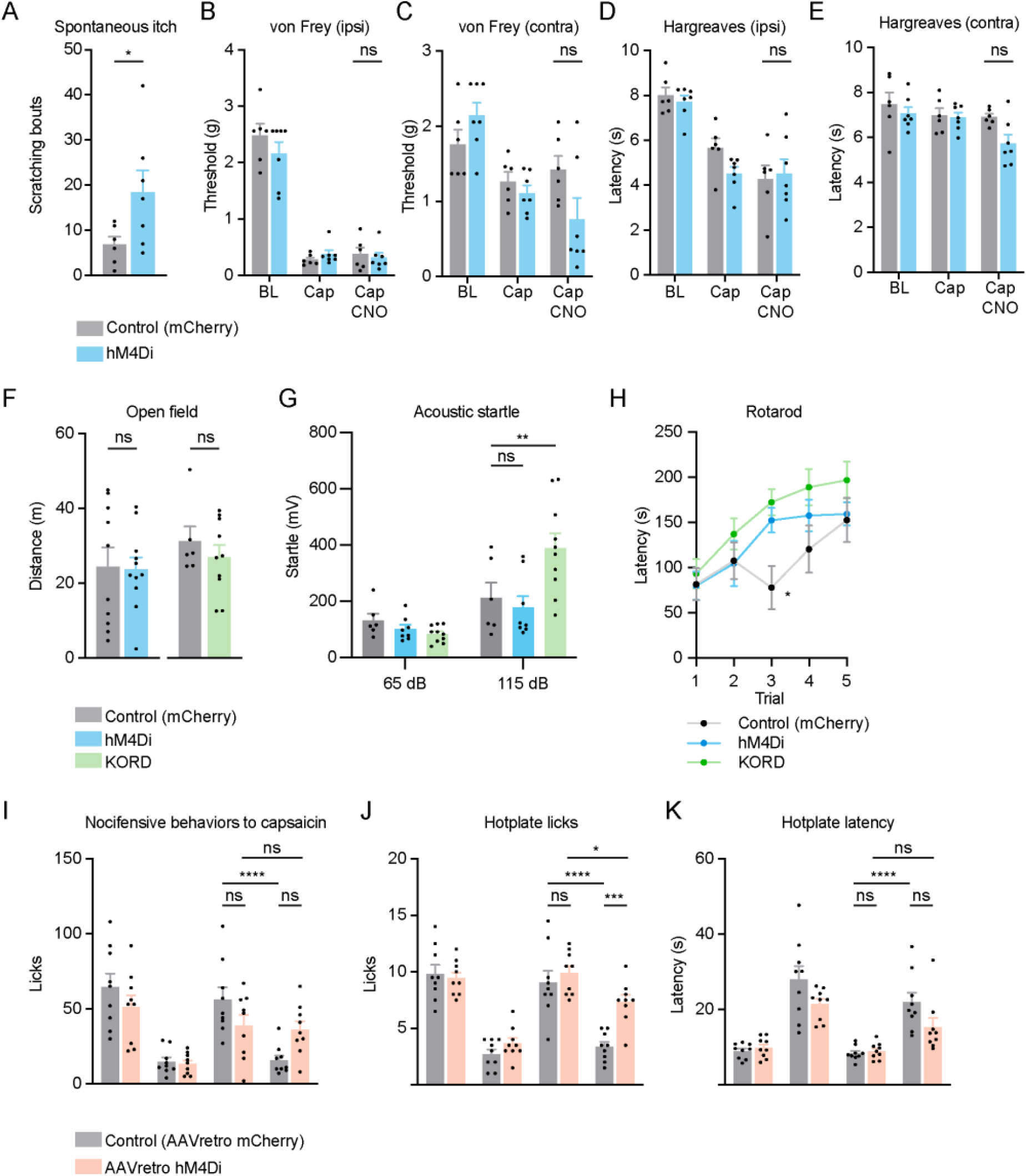
Supplementary chemogenetic inhibition behavioral data. **(A)** Effect of chemogenetic inhibition of RVM^KOR^ neurons on spontaneous itch. Data are mean + SEM with dots representing individual mice (n = 6 control, n = 7 hM4Di). Asterisks indicate the results of unpaired t-test (* p<0.05). **(B-E)** Effect of chemogenetic inhibition of RVM^KOR^ neurons on **(B, C)** mechanical and **(D, E)** thermal responses with capsaicin-induced injury. Data are mean + SEM with dots representing individual mice (n = 6 control, n = 7 hM4Di). NS indicates the results of two-way repeated measures ANOVA with Bonferroni’s correction (ns p>0.05). **(F)** Inhibition of RVM^KOR^ neurons with either hM4Di or KORD does not affect locomotor activity in an open field test. Data are mean + SEM with dots representing individual mice (n = 6 control, n = 8 hM4Di, n = 10 KORD mice). NS indicates the results of unpaired t-test (ns p>0.05). **(G)** Inhibition of RVM^KOR^ neurons with KORD, but not hM4Di enhances startle responses to acoustic stimulation at 115 dB. Data are mean + SEM with dots representing individual mice (n =6 control, n = 8 hM4Di, n = 10 KORD mice). Asterisks indicates the results of two-way ANOVA with Bonferroni’s correction (** p<0.01, ns p>0.05). **(H)** Inhibition of RVM^KOR^ neurons hM4Di and KORD generally did not affect performance on a rotarod. Although a difference was detected compared to the control group on Trial 3, this is likely due to a fluctuation in the performance of the control group. Data are mean ± SEM. n = 6 control, n = 8 hM4Di, n = 10 KORD mice. Asterisks indicates the results of two-way ANOVA with Bonferroni’s correction (* p<0.05). **(I-K)** Effect of chemogenetic inhibition of descending RVM^KOR^ neurons on stress-induced analgesia and responses to capsaicin-induced licking **(I)**, hotplate total licks **(J)** and latencies to lick on hotplate **(K)**. Data are mean + SEM with dots representing individual mice (n = 6 AAVretro control, n = 7 hAAVretro M4Di). Asterisks indicate the results of two-way repeated measures ANOVA with Bonferroni’s correction (*** p<0.001, **** p<0.0001, ns p>0.05).

**Supplementary Figure 5.**
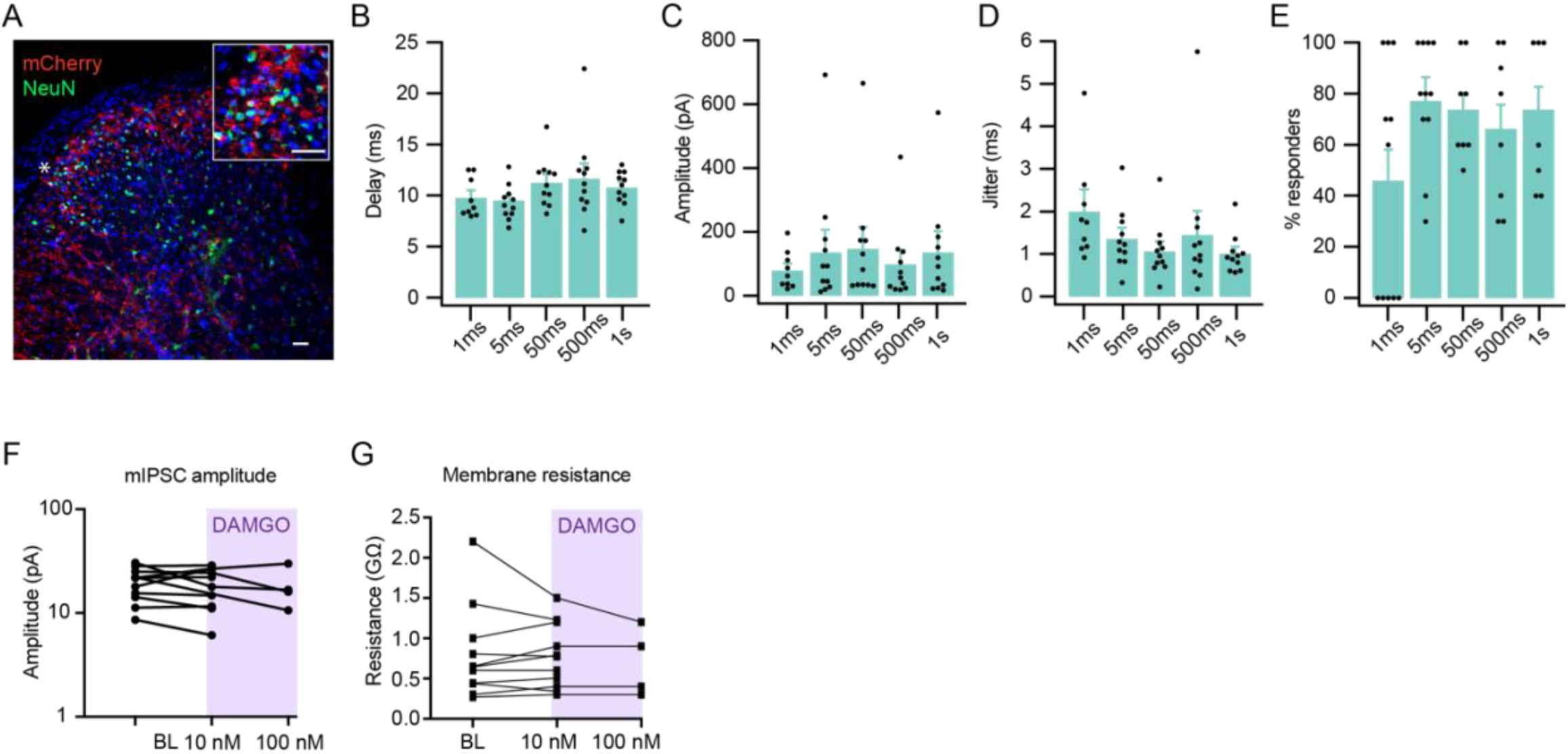
Supplementary spinal cord electrophysiology data. **(A)** Representative image of descending fibers originating from labeled RVM^KOR^ neurons in the spinal cord. Scale bar = 50 μm. **(B)** Delay between the optical activation of ChR2-expressing fibers in the spinal cord and IPSC onset in lamina II neurons at different durations of laser stimulation. (n = 11 cells from 8 animals). **(C)** Amplitude of IPSC at different durations of laser stimulation. (n = 11 cells from 8 animals). **(D)** Response jitter of IPSC at different durations of laser stimulation. (n = 11 cells from 8 animals). **(E)** Percentage of cells responding under different durations of laser stimulation. **(F-G)** Electrophysiological recordings of labeled, spinally-projecting RVM^KOR^ neurons before and after application of DAMGO (10 nM and 100 nM). Effects on mIPSC amplitude (F) and membrane resistance **(G)** are shown.

**Supplementary Figure 6.**
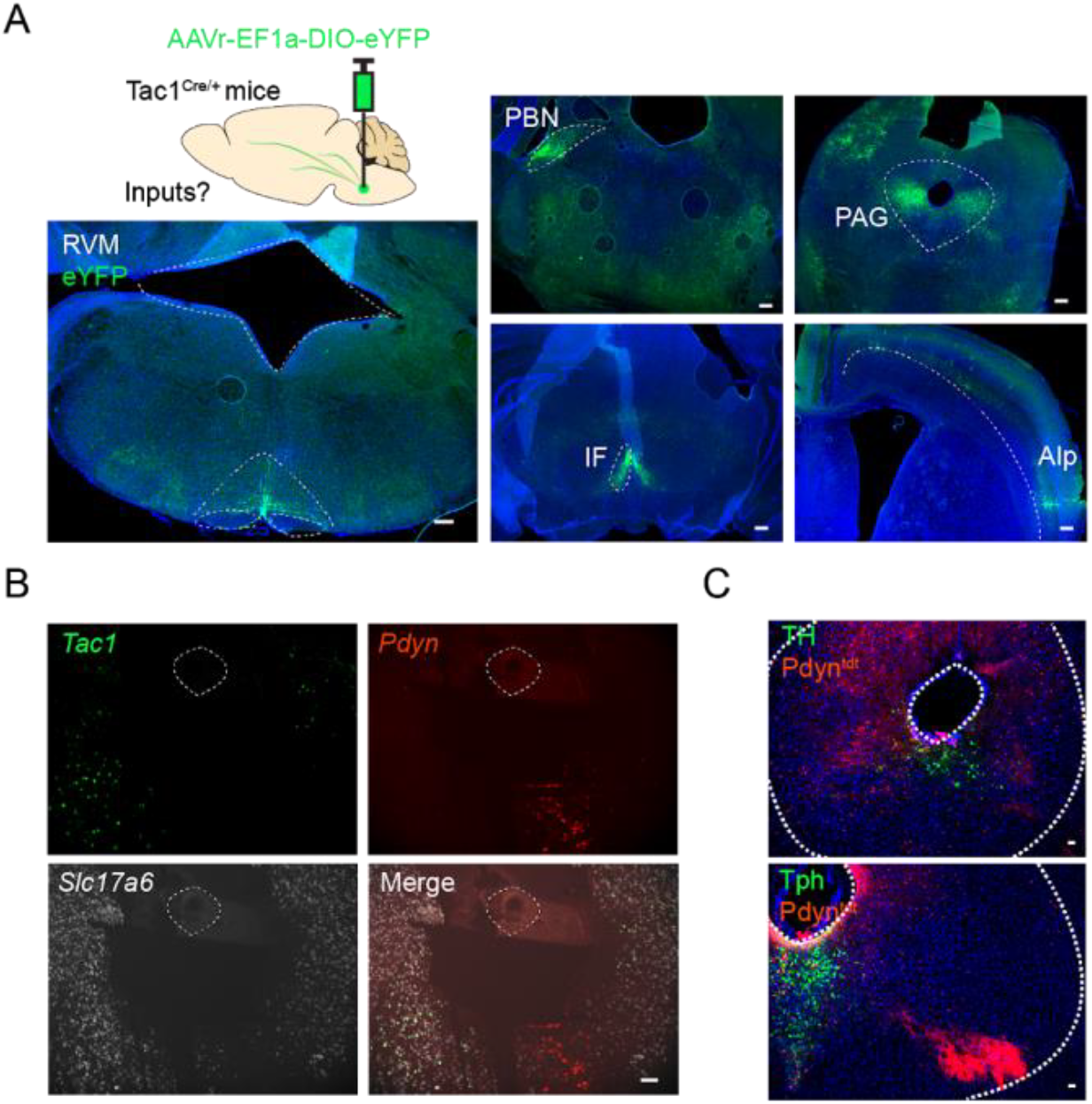
Further characterization of PAG neuronal cell types. **(A)** Experimental strategy to trace Tac1 inputs to the RVM. vlPAG: ventrolateral periaqueductal gray, PBN: parabrachial nucleus, AIp: agranular insular area, IF: interfascicular nucleus. Scale bar = 50 μm. **(B)** FISH characterization of *Pdyn, Tac1*, and *Slc32a1* expression in the PAG. Scale bar = 50 μm. **(C)** Immunohistochemical characterization of Dyn;tdt neurons and TH (tyrosine hydroxylase) and Tph (tryptophan hydroxylase). Scale bar = 50 μm.

